# MicroRNA Detection in Biological Media Using a Split Aptamer Platform

**DOI:** 10.1101/2022.01.17.476684

**Authors:** Liming Wang, Kern Hast, Tushar Aggarwal, Melih Baci, Jonathan Hong, Enver Cagri Izgu

**Affiliations:** Department of Chemistry and Chemical Biology, Rutgers University, New Brunswick, NJ 08854, USA; Cancer Pharmacology Program, Cancer Institute of New Jersey, Rutgers University, New Brunswick, NJ 08901, USA; Rutgers Center for Lipid Research, New Jersey Institute for Food, Nutrition, and Health, Rutgers University, New Brunswick, NJ 08901, USA

## Abstract

Intercellular microRNA (miRNA)-based communication has been implicated in a wide array of functional and dysfunctional biological processes. This has raised attention to the potential use of miRNAs as biomarkers for disease diagnosis and prognosis and produced interest in their detection. Though the list of clinically significant miRNA biomarkers is rapidly expanding, it remains challenging to adapt current tools to investigate new targets in biological environments. Systematic approaches for the rapid development of miRNA biosensors are valuable to reduce this disparity. We describe here a methodology for developing aptamer-based fluorescent biosensors that can specifically detect miRNAs in biological environments, including culture medium from HeLa cells, human serum, and human plasma. This methodology includes the semi-rational design of the hybridization between a pair of split DNA aptamer oligonucleotides and the miRNA target to build a pool of potential sensor designs, and the screening of this pool for designs with high signal-to-background ratio and sequence selectivity. The method uses natural oligonucleotides without chemical modification, and is effective in buffer, 10%, and 30% (v/v) biological media. Following this approach, we developed sensors that detect three miRNA targets (miR-19b, miR-21, and miR-92a) at concentrations as low as 5 nM without amplification and are selective against single-nucleotide mutants. This work expands upon the current design principles of nucleic acid-based biosensors and provides a method to rapidly develop diagnostic tools for novel and niche miRNA targets of interest.

## INTRODUCTION

MicroRNAs (miRNAs) are ca. 22 nucleotide-long noncoding RNA oligomers that contribute to post-transcriptional regulation of gene expression, typically via facilitation of degradation of complementary messenger RNAs in a process known as RNA silencing^1^. Recently, intercellular communication employing miRNAs as signaling molecules has acquired much attention^2^ due to its role in the regulation of several biological processes, including cell homeostasis, cell fate specification, and embryonic development^3,4^. When the dysregulation of these processes results in disease, miRNAs can serve as important disease markers. Tumor cells secrete miRNAs to facilitate cancer progression and metastasis by altering gene expression in neighboring cells^5,6^. The bodily fluids of cancer patients typically show elevated concentrations of certain miRNAs^7^. This has suggested the potential for their use as biomarkers for cancer diagnosis and prognosis^8–10^ and stimulated recent developments in miRNA diagnostics ^11–14^.

As the critical roles of miRNAs continue to be illuminated, there is a growing need for investigative platforms that are easily accessed and configurable for the detection of different miRNA targets in biological environments. However, the sequence space of the miRNAs is large. The human transcriptome alone contains over 2,600 miRNA sequences and over 48,000 have been identified in all organisms studied^15^. Independent approaches for the development of unique tools for each miRNA sequence is inadequate to address this large sequence space. Systematic approaches for cost-effectively and rapidly producing tools for sequence-specific miRNA detection can greatly increase the accessibility of studying miRNA of interest.

One technology with significant potential for RNA detection and cancer diagnosis ^16^ is the aptamer-based biosensor. Aptamers, which are oligonucleotides that exhibit high binding affinity toward a specific ligand ^17^, can produce fluorescence without covalent labeling and can be reconstituted to specifically hybridize with a nucleic acid target^18^. Furthermore, the ability to function at room temperature provides an advantage over molecular beacon sensors that require elevated temperatures to differentiate nucleic acid mutants^19^. A promising aptamer-based nucleic acid-sensing platform^20^ is composed of a split form of the malachite green aptamer (MGA), an *in vitro*-selected, light-up RNA aptamer to which malachite green (MG) tightly binds^21^ and enhances its fluorescence^22^. In this split aptamer design, MGA was separated into two strands that must come together to form the MG-binding motif. Each aptamer strand contains a 7-nucleotide (nt) region complementary to a 14-nt target sequence. Hybridization of the aptamer strands with a single-stranded DNA oligomer containing the target sequence stabilizes the folding of the MG-binding motif, facilitating binding of MG and thus increasing its quantum yield.

Despite their potential, aptamer-based nucleic acid detection tools have yet to meet certain requirements for practical application as miRNA biosensors. First, the system must hybridize only with specific sequences of ca. 22-nt, ensuring that the sensor is specific to the miRNA target against the vast sequence space found in transcriptome. This is challenging, because as the length of the hybridizing region increases, so too does the strength of the hybridization and the tolerance of mismatches between the sequences. Secondly, these biosensors must operate in the complex environments native to cells and tissues. These environments contain a multitude of confounding factors, including non-target oligonucleotides, nucleases that can degrade the aptamer, and a cocktail of biomolecules that may interfere with the ligand-aptamer interaction. Most reported assays are performed in either buffers or highly diluted biological samples. Finally, because of the vast number of potential miRNA targets, it is critical to design the aptamer platform with minimal or preferably no chemical modification to allow accessibility and configurability. Methods that can produce a wide array of miRNA sensors that overcome these challenges are needed.

Here we report a systematic methodology for producing split aptamer biosensors that can detect miRNA targets in various biological media, including culture medium collected from HeLa cells (CM), human serum (HS), and human plasma (HP). The biosensors produced from this method require minimal sample handling and can detect miRNA with nanomolar levels of sensitivity and single-nucleotide specificity. The split aptamer strands are composed of DNA, providing the system with improved stability against degradation. Furthermore, these strands use the natural polymer of DNA, without chemical modification, rendering the production of novel biosensor designs rapidly accessible and cost effective.

Our platform, inspired by the split MGA system^20^, utilizes a fluorophore binding region that is stabilized upon hybridization with the miRNA target. This simple approach provides significant tunability to the structure of the hybridization region, and our method of biosensor development focuses on the optimization of the miRNA-interacting region to maximize stability and signal generation in the presence of the miRNA target and minimize stability and background signal in the absence of it. Our method has three steps: 1) use semi-rational design to construct a pool of potential biosensors with diverse structural characteristics, 2) screen this pool against a specific miRNA target to identify designs with high signal-to-background ratios, and 3) screen the identified designs against targets with single-nucleotide mutations to select complexes with high sequence specificity. The semi-rational design of potential biosensors narrows the sequence space to a manageable list, of which further screening can robustly and rapidly produce effective sensors.

We show the flexibility of this methodology by designing biosensors for three miRNA targets with relevance to dysfunctional cellular physiology: miR-19b, miR-21, and miR-92a. MiR-19b and miR-92a are part of the miR-17/92 cluster—a group of related miRNA genes that plays a regulatory role in the metabolic reprogramming of lymphoma cells and promotes Myc-dependent tumor growth^23^. Consistently elevated levels of these two miRNAs are correlated with silencing of critical tumor suppressor genes^23, 24^. The third target, miR-21, downregulates tumor-suppressing apoptotic proteins, and its overexpression is observed in breast, ovarian, colorectal, gastric, prostate, and lung cancers^25^. All three of these miRNAs were detected in the bodily fluid of patients with diffuse B-cell lymphoma at mean expression levels 10 to 16-fold higher than that of healthy individuals^7^. Important to the development of aptamer-based biosensors, these miRNA targets all have different chemical features that challenge the design of a sensor vis-à-vis secondary structures and hybridization.

All three sensors developed by this methodology displayed good single-nucleotide specificity for their targets in both buffer and biological media. At 2 μM miRNA concentration, an ca. 20-fold signal enhancement in both buffer and 10% (v/v) HS, and a 10-fold signal enhancement in 30% (v/v) HS was reached. Without amplification, the sensors could detect miRNAs at low nanomolar levels at room temperature within one hour in both buffer and the biological media.

## RESULTS AND DISCUSSION

### Split Aptamer Design

As a template for our aptamer designs, we devised a multi-stranded complex (**Fig. 1**) formed from two DNA strands (St1 and St2) and the target miRNA strand. The miRNA sequences (**Supplementary Table 1**) were analyzed with the Vienna RNA websuite^26^ for secondary structure and hybridization. MiR-19b has a strong secondary structure, miR-21 can self-dimerize, and miR-92a has the highest hybridization energy, including a 7-nt stretch of C and G nucleotides (**Supplementary Fig. 1**). The St1:St2:miRNA complex possesses a ligand-binding region (LigBR) and four stems that stabilize the folding of this region. The LigBR is formed from the two DNA strands and is a consensus sequence to which a small-molecule fluorophore binds. For the LigBR, we utilized the sequences 5′-GGGGGAGGGTGTGTGGTCTT-3′ and 5′-GCTTGGTTC-3′, which form a split version of an *in vitro*-selected motif to which dapoxyl dyes fluorogenically bind, termed DAP-10 ^27^. However, dapoxyl-family molecules have been reported to fluorogenically bind human serum albumin^28^, which limits their application in clinical assays. In agreement with this report, our tests showed that dapoxyl sulfonate, a dye previously used with DAP-10 ^27^, exhibited strong fluorescence without DAP-10 in HS, HP, cell media containing 10% (v/v) fetal bovine serum, and in buffers containing 1% (v/v) fetal bovine serum or 25 μg/mL bovine serum albumin (**Supplementary Fig. 2a-b**). We also observed that dapoxyl sulfonyl fluoride, an analog of dapoxyl sulfonate that has recently been used with split aptamers to detect gene fragments of bacterial and viral pathogens in buffer^29^, showed intense non-specific fluorescence in biological media. In our hands, dapoxyl sulfonyl fluoride had only 2-fold fluorescence enhancement when mixed with DAP-10 in a fluorescence assay buffer (FAB: pH 7.6, 20 mM Tris, 140 mM NaCl, 5 mM KCl, and 2 mM MgCl2), but without DAP-10 in biological media (10% v/v) generated a 9 to 17-fold increase over the intrinsic fluorescence of the media (**Supplementary Fig. 2c**). Another small-molecule dye, Auramine O (AO), was recently reported to generate strong fluorescence when mixed with DAP-10^30^ and notably was used to study mycobacteria-induced neutrophil phagocytosis in human plasma^31^. In our investigations, fluorescence enhancement of AO without DAP-10 was minor in biological media (2 to 10-fold), whereas that with DAP-10 was significant in both FAB (3,800-fold) and biological media (82 to 494-fold, **Supplementary Fig. 2d**). Due to its superior performance in biological media over the dapoxyl family dyes, we chose AO as the aptamer ligand for building our biosensors.

**Figure 1.**
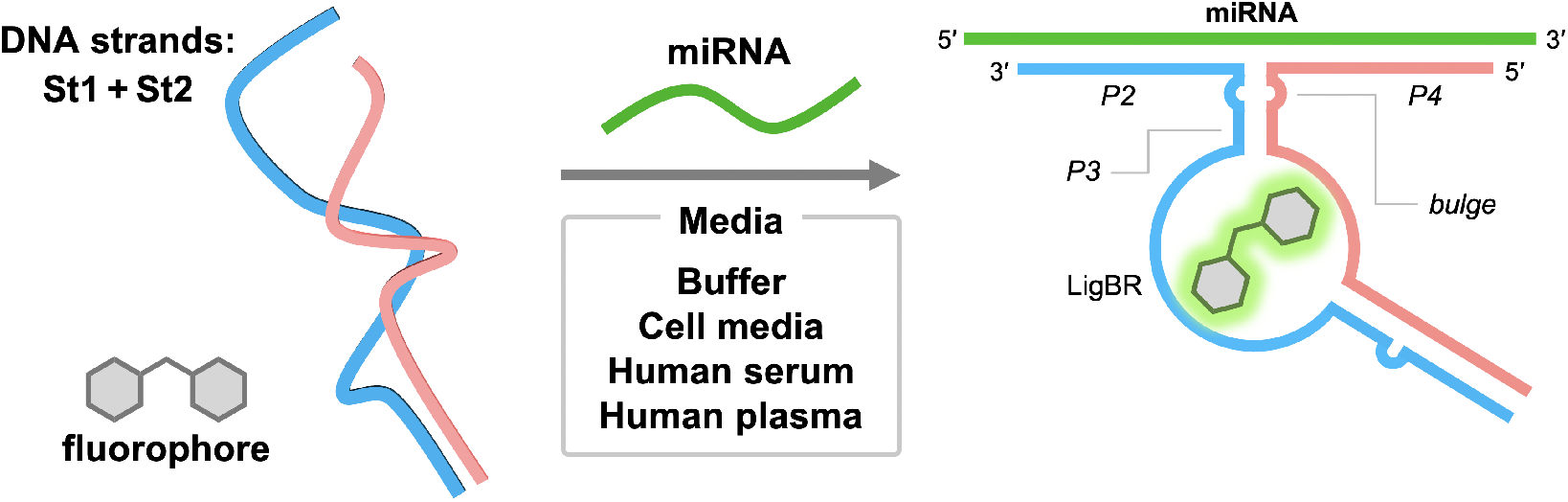
The working principle of miRNA detection via split aptamer folding and fluorophore binding. In the absence of the target miRNA, the ligand binding region (LigBR) is destabilized and has low affinity for the fluorophore. When the miRNA is added, its hybridization with St1 and St2 stabilize the LigBR, increasing its affinity for the fluorophore, the binding of which increases the fluorescent output of the fluorophore.

**Figure 2.**
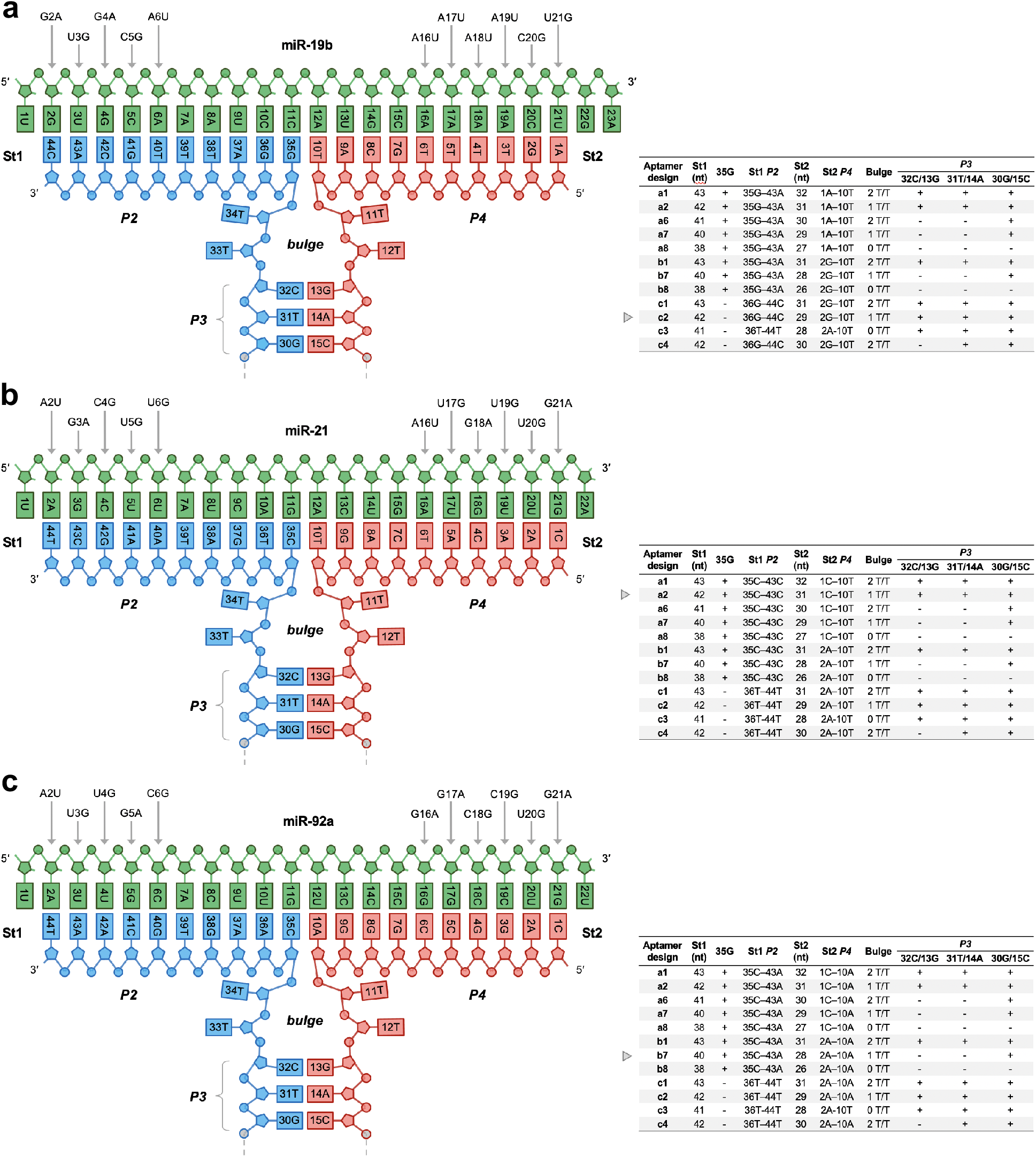
Sequence and secondary structures of the highest performing aptamers targeting (a) miR-19b, (b) miR-21, and (c) miR-92a. Single nucleotide mutations are denoted with an arrow above each miRNA sequence. Parameters used for each aptamer design are listed on the right, with the drawn aptamer denoted by a triangle.

In our semi-rational design, stems P1 and P3 are formed on either side of the LigBR by the DNA strands and stabilize folding of the LigBR. P3, which closes the LigBR on the side nearest to the miRNA-binding region, is composed of a short stretch of base pairs, which stabilizes LigBR upon formation of P2 and P4 in the presence of the target miRNA. Stems P2 and P4 are each formed between a DNA strand and the 5′ and 3′ regions of a miRNA, respectively, and form a Y-junction with P3. In the presence of the target miRNA strand, the formation of P2 and P4 stabilizes the formation of P3, which in turn facilitate the proper folding of the LigBR. By adjusting the lengths of P2, P3, P4, and the flexibility of the Y-junction formed by these 3 stems, we can manipulate the stability and folding of the LigBR in the presence and absence of the miRNA. The lengths of the stems affect their hybridization strengths. Complementary sequences with seven or more contiguous base pairs typically have favorable annealing rates and duplex stability in physiologically relevant conditions^32^. Building on this, we sought to install P2 and P4 stems that are 8 to 10-bp long—reasoning that in this range even a single nucleotide mutation in the miRNA sequence should result in a considerable decrease in stem stability. As P3 must be destabilized in the absence of the miRNA, a shorter stem is required, and thus we produced designs where P3 was 0 to 3-bp long. Additionally, the steric effects of the three helices converging at the Y-junction may produce significant strain on the formation of the stems and LigBR, so the flexibility of the Y-junction was tuned by the following two parameters: 1) relaxing effect of a bulge 0 to 2 nt long at the junction on either side of P3, 2) an unpaired 11N in the miRNA at the Y-junction. Using combinations of these five parameters, a pool of variations on the split aptamer template can be screened for high-performing designs. While it is reasonable that these parameters are likely to be important across all miRNA targets, it should be noted that the design that is most optimal for each miRNA may be different depending on the miRNA sequence and structure.

Implementing these principles, we first screened a pool of 40 different St1:St2 sets with different structural characteristics (**Supplementary Table 2** and **3**). The St1:St2 set names are given a letter and number designation and suffixed with the miRNA they target, e.g., set a1-92a refers to St1:St2 design a1 configured to complex with miR-92a (See **Supplementary Table 2** for a complete list of designations with meanings of the letter and number designations). These St1:St2 sets were then tested with miR-19b to identify the systems with the highest fluorescence signal-to-background ratios, *F/F_0_*, where *F* and *F_0_* are the fluorescence intensities of the assay with and without the target miRNA, respectively. A larger *F/F_0_* indicates a better signal to noise ratio. Through this screening, we identified twelve St1:St2 sets (a1-19b, a2-19b, a6-19b, a7-19b, a8-19b, b1-19b, b2-19b, b7-19b, b8-19b, c1-19b, c2-19b, c3-19b, and c4-19b; see **Fig. 2a**) that each exhibited an *F/F_0_* of over 5.

We then evaluated the sequence specificity of these twelve sets by comparing their fluorescent output with native miR-19b to that with each of a set of eleven miR-19b single-nucleotide mutants. A differentiation factor, *D_f_*, was calculated according to the formula

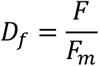

where *F* is the fluorescence intensity of the assay with native miR-19b *and F_m_* is the fluorescence intensity with the miR-19b mutant. A *D_f_* of 1 indicates no specificity for either sequence, while a larger *D_f_* indicates the fold increase in specificity for the native miRNA sequence. For many St1:St2 sets, *D_f_* for all the mutants was larger than 1, with many mutants showing a *D_f_* in the range of 3-10 (**Fig. 3b**). For miR-19b, *D_f_* typically increased when the mutation was located closer to the 3′-end of P2 or 5′-end of P4 (*i.e*., far from LigBR), suggesting that pairings at these ends had less impact on the folding of the consensus region.

### Configurability for General MiRNA Detection

To show the generalizability of this approach to other miRNA targets, we changed the DNA sequences in the P2 and P4 stems of the selected group of twelve St1:St2 sets to complex with two other miRNA targets, namely miR-21 (**Fig. 2b**) and miR-92a (**Fig. 2c**). *F/F_0_* and *D_f_* for each new St1:St2:miRNA set were measured using the same fluorescence assay protocols described above. For miR-21, an *F/F_0_* > 5 was observed for assays with a2-21, a6-21, a7-21, c2-21, and c4-21 (**Fig. 3d**). A *D_f_* larger than 3 was also observed for these five sets (**Fig. 3e**). For miR-92a, the *F/F_0_* of all complexes except a1-92a, a2-92a, b1-92a, and c3-92a were 10 or higher (**Fig. 3g**) and *D_f_* over 3 was observed for most miR-92a mutants (**Fig. 3h**). Notably, the G16A mutant of miR-92a had a *D_f_* less than 2 across all the St1:St2 sets. This suggests that this mismatch, which is positioned at the middle of the 7-nt G and C segment of miR-92a (13C–19C), is adequately stabilized by the flanking, strongly hybridized C/G pairs.

**Figure 3.**
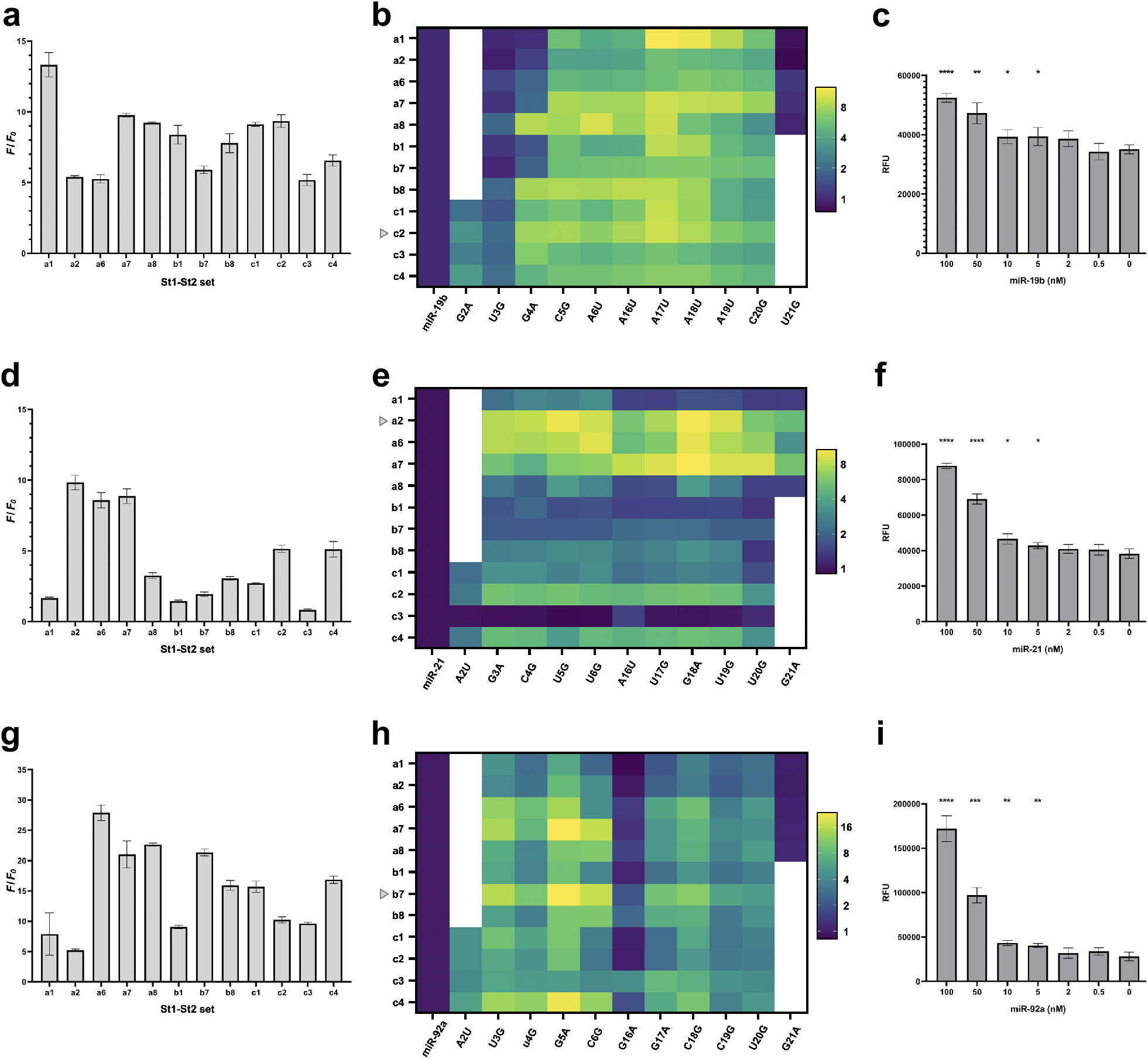
Performance of the split aptamer platform in fluorescence assay buffer (FAB). (**a**, **d**, and **g**) The fold enhancement of the fluorescent signal with the addition of miR-19b, miR-21, and miR-92a, respectively, for the high performing complexes. (**b**, **e**, and **h**) The fold enhancement of the fluorescent signal with the addition of single nucleotide mutants of miR-19b, miR-21, and miR-92a, respectively. The designs used for biological media testing are denoted with a triangle. A log2 scale is used for coloration. (**c**, **f**, and **i**) LoD measurements in FAB for c2-19b, a2-21, and b7-92a, respectively. Error bars represent the standard deviation; n = 3. Single-tailed Student’s *t* test: *, P < 0.05; **, P < 0.01; ***, P < 0.001; ****, P < 0.0001.

For each miRNA target, one high-performing complex, namely c2-19b, a2-21, and b7-92a, was selected for further characterization and testing in biological media.

### Ligand Affinity of the Split Aptamer with and without Target miRNA

The dissociation constant of AO bound to the LigBR, *K_d_*, for each of the three high-performing complexes was determined by fluorescence assay. Different concentrations of each complex were incubated with 1 μM AO until the fluorescence plateaus, and the fluorescence intensities were fitted to a 1:1 binding model. The *K_d_* for AO binding with the miRNA was based on curve fitting, whereas that without miRNA was estimated based on *F/F_0_*, as the binding affinities were too weak to calculate *K_d_* at practical concentrations of St1 and St2 strands. The *K_d_* for AO with c2-19b:miR-19b complex was 9.1 μM (**Supplementary Fig. 4a**), while the *K_d_* for AO with only c2-19b was 180 μM (**Supplementary Table 4**). Next, the *K_d_* for AO with a2-21:miR-21 was 4.6 μM (**Supplementary Fig. 4b**), while the *K_d_* for AO with a2-21 was 140 μM based on *F/F_0_*. Finally, the *K_d_* for AO with b7-92a:miR-92a was 4.2 μM (**Supplementary Fig. 4c**), while the *K_d_* for AO with b7-92a was 310 μM. The significant increase in *K_d_* in the absence of the target miRNA supports the proposed mechanism of our split aptamer design wherein the presence of the target miRNA strengthens the interaction of AO and LigBR.

**Figure 4.**
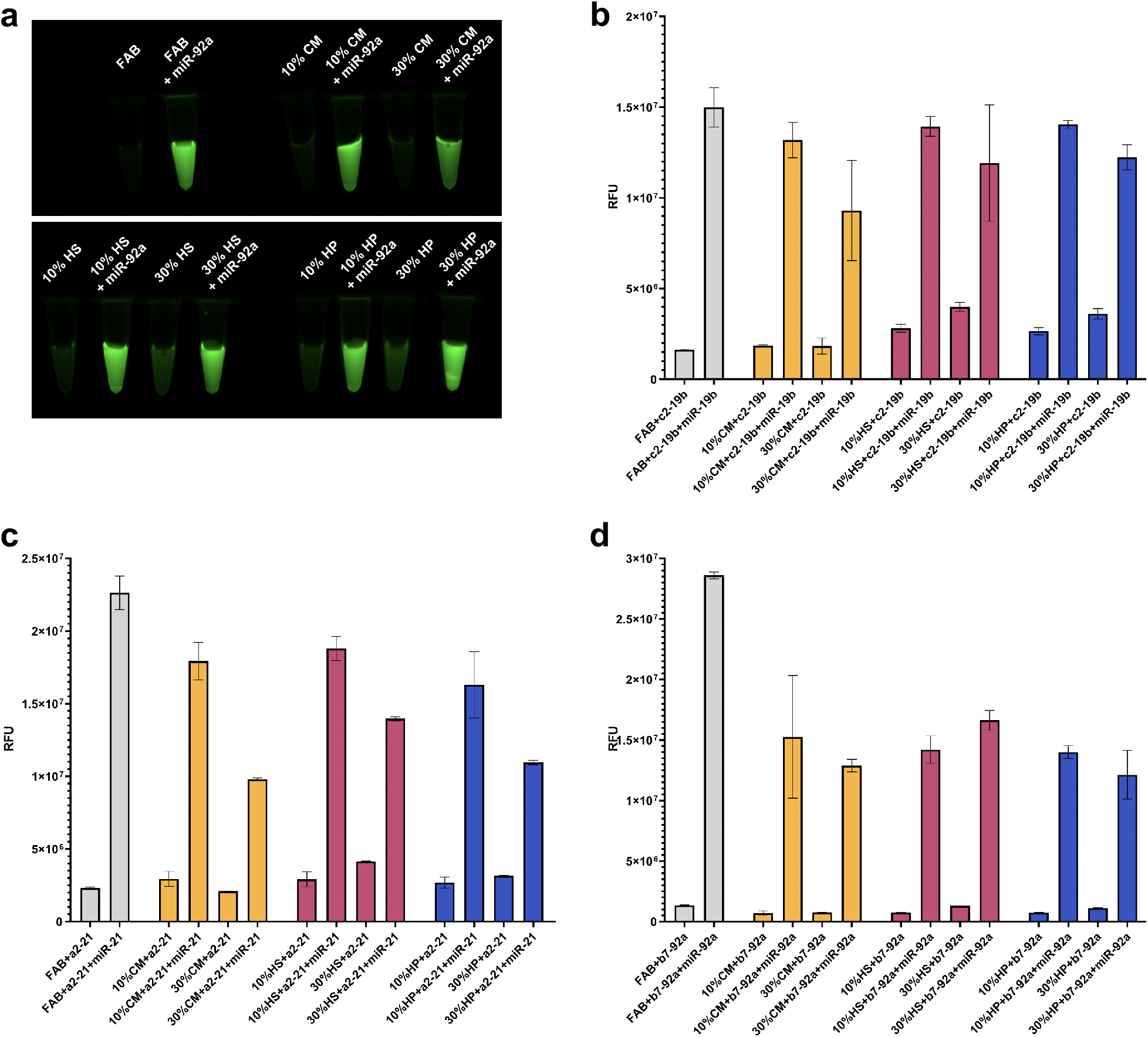
Target sensitivity of the binary aptamer platform in different media. (**a**) Fluorescence imaging of miR-92a using b7-92a in fluorescence assay buffer (FAB), cell media (CM), human serum (HS), and human plasma (HP). (**b-d**) Fluorescence intensities of the three selected split aptamer systems with their target miRNA. Error bars represent the standard deviation; n = 3.

### Limit of Detection (LoD) in FAB

We determined the LoDs for the high performing complexes by assessing the statistical difference between *F* and *F_0_* at various concentrations of the target miRNA. Here, LoD is defined as the minimum miRNA concentration at which *F* is statistically different than *F_0_* using the one-tailed Student’s *t* test. We found that the sensitivities were within an order of magnitude for each high performing design, all presenting an LoD of 5 nM in FAB (**Fig. 3c**, **3f**, and **3i**). These sensitivities are comparable to those displayed by other nucleic acid probes that do not utilize amplification, such as molecular beacons^19^, catalytic assemblies of LigBRs^33^, and modified oligonucleotides that ligate to a target sequence^34^.

### MiRNA Detection in Biological Media

We evaluated the application of the aptamer biosensor to miRNA detection in complex biological environments, such as those present in cell culture and clinical samples (**Fig. 4**). We first prepared 10% (v/v) and 30% (v/v) solutions of three different media in FAB, namely heparinized human plasma, human serum clotted from whole blood, and media from cultured HeLa cells. It should be noted that heparinized plasma was chosen over plasma with anti-coagulants other than heparin, such as EDTA and citrate, as they can chelate the magnesium ions in the buffer, which are important for aptamer activity. These media solutions were spiked with 2 μM of the appropriate miRNA target, and murine RNase inhibitor was added to a final concentration of 1 unit/μL to stop degradation of the spiked miRNA. *F/F_0_* for each of the three high-performing complexes were measured using the same fluorescence assay protocols described above. The *F/F_0_* for c2-19b with miR-19b in 10% (v/v) CM, HS, and HP were 7.1, 5.0, and 5.3, respectively; and in 30% (v/v) CM, HS, and HP were 5.1, 3.0, and 3.4, respectively (**Fig. 4b**). The *F/F_0_* for a2-21:miR-21 in CM, HS, and HP were 7.2, 6.2, and 5.5, respectively; and in 30% (v/v) CM, HS, and HP were 4.7, 3.4, and 3.5, respectively (**Fig. 4c**). Finally, the *F/F_0_* for b7-92a:miR-92a in CM, HS, and HP were 21.8, 19.2, and 19.2, respectively; and in 30% (v/v) CM, HS, and HP were 17.1, 12.8, and 10.8, respectively (**Fig. 4d**). The signal-to-background ratios of all the three sensors were retained in the biological media, and the three split aptamer designs showed satisfactory performances in both FAB and biological media.

Next, we tested the split aptamer platform with the miRNA mutants that displayed both the highest and lowest *D_f_* in FAB to assess whether it retains sequence specificity in biological media. *D_f_* was measured using the same fluorescence assay protocol described above (**Supplementary Fig. 5a-c**). At 2 μM concentration, the miR-19b mutant A17U (*D_f_* = 9.8 in FAB) exhibited similar signal intensities across all the assay media, with *D_f_* values for CM, HS, and HP centered around 11.1 (**Supplementary Table 5**). For the mutant U3G (*D_f_* = 2.0 in FAB), sequence differentiation was also satisfactory, with *D_f_* values ranging from 2.8 (in HeLa cell media) to 14.3 (in human serum). In the case of miR-21 and miR-92a mutants, sequence differentiation was retained as well. The miR-21 mutant G18A (*D_f_* = 10.2 in FAB) resulted in similar *D_f_* in biological media, ranging from 6.7 to 12.5, while for mutant G21A (*D_f_* = 5.1 in FAB), the *D_f_* in biological media centered around 7.7. As for miR-92a, mutant G5A (*D_f_* = 23.8 in FAB) presented a *D_f_* around 14.3, and the sequence differentiation of mutant G16A (*D_f_* = 2.0 in FAB) was slightly improved, presenting *D_f_* ranging from 3.2 to 5.0 in biological media. Taken together, these results indicate that the split aptamer platform performs with a considerable level of sequence specificity in biological media, even at the 30% media concentration.

**Figure 5.**
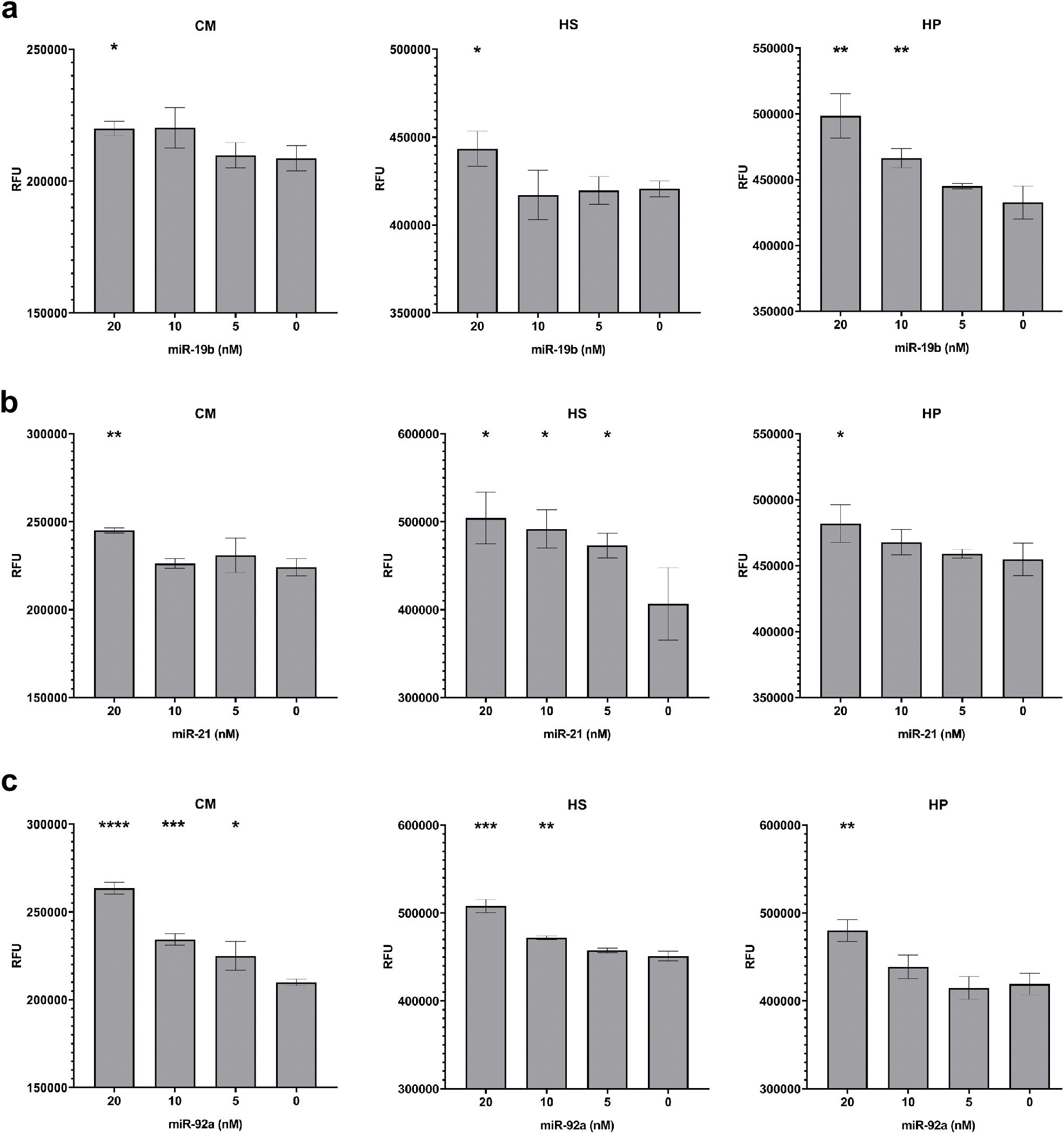
LoD measurements for (**a**) miR-19a, (**b**) miR-21, and (**c**) miR-92a in biological media (10% v/v). Error bars represent the standard deviation; n = 3. Single-tailed Student’s *t* test: *, P < 0.05; **, P < 0.01; ***, P < 0.001; ****, P < 0.0001.

### LoD in Biological Media

To further assess whether the sensitivity of our selected designed aptamer platforms is retained in biological media, we determined the LoDs in the three biological media at 10% v/v. The LoDs for all assays displayed in a range between 5 and 20 nM in the three biological environments (**Fig. 5**). Remarkably, the LoDs of a2-21 in HS and b7-92a in CM were determined as 5 nM, the same as their LoD in FAB, indicating that their performance was retained in these environments very well. The LoDs of our selected sensors in three biological media were close to those in FAB, suggesting that the sensitivity of these platforms was comparable in biological environments to the sensitivity in FAB.

## CONCLUSION

Accurate detection of specific miRNAs remains challenging due in part to the sequence similarity in the transcriptome and other confounding factors within biological environments, such as background interference and biochemical degradation.

In this work, we presented a methodology that includes the semi-rational design of DNA strands hybridizing with a miRNA target to generate a pool of potential miRNA sensors and the screening of this pool to identify those with high signal-to-background ratio and sequence selectivity. The reported modular and easily employed split aptamer platform accurately detected target miRNAs in the presence of complex biological content and near matches. The adaptability of this platform was demonstrated by designing sensors for three oncogenic miRNAs: miR-19b, miR-21, and miR-92a. Without amplification, these miRNAs were detected at low nanomolar levels at room temperature within one hour. The two backbone strands of the split aptamer design were built from DNA polymers without chemical modification.

This work expands the chemical repertoire available for sensing cancer biomarkers by allowing detection within the environments native to cells and tissues. The biosensing platform described here has potential for improving insight into miRNA-based intercellular communication pertinent to disease progression. The ability to attain fidelity at the single nucleotide level opens the possibility of identifying potential genetic or epigenetic cancer biomarkers with differences that affect base pairing.

## ACKNOWLEDGMENTS

This work was supported by the US National Institutes of Health / NIBIB Trailblazer Award (R21 EB029548), American Cancer Society, Institutional Research Grant Early Investigator Award, and the Rutgers Cancer Institute of New Jersey NCI Cancer Center Support Grant (P30CA072720). We acknowledge Dr. M. Rhia L. Stone for preparation of HeLa cell media. We thank Prof. Zheng Shi and Shilong Yang for providing us with the HeLa cell line, and Dr. Venu Gopal Vandavasi for helpful discussions.

## AUTHOR CONTRIBUTIONS

E.C.I. conceived the study. L.W. and T.A. conducted biosensor design and screening, L.W. and K.H. performed biological assays and thermodynamic analyses, M.B. synthesized small-molecule fluorophores, J.H. identified clinically and chemically relevant miRNA targets. All authors contributed to the interpretation of data. L.W., K.H., and E.C.I. wrote the manuscript with input from all authors. E.C.I. supervised the study.

