## Supplementary Information for "MicroRNA Detection in Biological Media Using a Split Aptamer Platform"

| **Contents** | **Page** |
|---|---|

| | |
|---|---|
| **Chemicals and Materials** | S3 |
| **General Synthetic Methods** | S4 |
| **Preparation Procedures and Characterization Data for DSF and DS** | S5–S13 |
| **DSF** | S5 |
| **DS** | S6 |
| Copies of Spectral Data ($^{1}$H, $^{13}$C, $^{19}$F NMR, and MS) | S7–S13 |
| **Preparation and $^{1}$H NMR Quantification of AO** | S14–S15 |
| **MiRNA Secondary Structures and Hybridization Energies** | S16 |
| **Comparative Analysis of Fluorogenicity of DSF, DS, and AO in Buffer and Biological Media** | S17 |
| **MiRNAs and MiRNA Mutants** | S18–S19 |
| **Split Aptamer Design** | S19–S21 |
| **MiRNA Detection** | S22 |
| **Dissociation Constant Determination** | S23–S24 |
| **Sequence Specificity in Biological Media** | S25–S27 |
| **Limit of Detection (LoD)** | S28 |
| **References** | S28 |

### Chemicals and Materials

**Reagents:** 2-Amino-4'-dimethylaminoacetophenone hydrochloride was purchased from Combi Blocks. Pyridine, methylene chloride, chloroform, sodium hydroxide, sodium carbonate (all anhydrous), HPLC grade acetonitrile, ACS grade acetone, 4-(fluorosulfonyl)benzoyl chloride, auramine O (AO, 85% dye content), benzoic acid, and bovine serum albumin (BSA), were purchased from Millipore-Sigma. 98% sulfuric acid was purchased from VWR Chemicals BDH. 2-amino-4'-dimethylaminoacetophenone was purchased from TCI America. Dulbecco's Modified Eagle Medium (DMEM) with phenol red was purchased from Corning. 10% (v/v) fetal bovine serum (FBS) was purchased from Hyclone Laboratories Inc. Dapoxyl sulfonate (DS) and dapoxyl sulfonyl fluoride (DSF) were synthesized according to published procedures[1] with modification (detailed below).

**Buffers and Solvents:** Tris(hydroxymethyl)aminomethane (Tris) was purchased from Millipore-Sigma. Water was deionized and filtered to a resistivity of 18.2 ΩM with a Milli-Q® Plus water purification system (Millipore, Massachusetts). Buffers were prepared freshly in Milli-Q® water and their pH was adjusted using HCl or NaOH. Fluorescence assay buffer (FAB): 20 mM Tris, 140 mM NaCl, 5 mM KCl, 2 mM $MgCl_2$, pH 7.6.

**Biological Media:** CM: Cell media, HS: Human serum, and HP: Human plasma.
CM was obtained from HeLa cell (CCL-2, ATCC) cultured in DMEM that was supplemented with 10% FBS. HS was purchased from Millipore-Sigma as heat inactivated, from male AB clotted whole blood. HP was purchased from Equitech-Bio, Inc. as human unfiltered sodium heparin plasma.

**Oligonucleotides:** DNA and RNA sequences used in this work were obtained as dry thin film from Millipore-Sigma. All of the oligonucleotides were cartridge purified by the manufacturer. Stock solutions of oligonucleotides were prepared with Milli-Q® Plus water and stored at -80 °C.

### General Synthetic Methods

For the chemical synthesis of organic compounds, all reactions were performed under a dry nitrogen atmosphere unless otherwise stated. All glassware was oven-dried before use. Purification of the synthesized compounds was performed using a Büchi Reveleris® flash chromatography system equipped with a FlashPure EcoFlex C-18 (50 μm sphere) column. Nuclear Magnetic Resonance (NMR) spectroscopic analyses were carried out on either a Bruker Avance Neo 500 MHz or Varian VNMRS 500 MHz spectrometer. $^{1}$H NMR spectra were acquired at 500 MHz, $^{13}$C NMR spectra were acquired at 126 MHz, and $^{19}$F NMR spectrum was acquired at 471 MHz. Chemical shifts ($\delta$) for $^{1}$H NMR spectra were referenced to $(C\boldsymbol{H}_3)_4Si$ at $\delta$ = 0.00 ppm and to $C\boldsymbol{H}D_2S(O)CD_3$ at $\delta$ = 2.50 ppm. $^{13}$C NMR spectra were referenced to $\boldsymbol{C}D_3S(O)\boldsymbol{C}D_3$ at $\delta$ = 39.52 ppm. The following abbreviations are used to describe NMR resonances: s (singlet) and d (doublet). Coupling constants (*J*) are reported in Hz. Liquid chromatography followed by high-resolution mass spectroscopy (LC-HRMS) analysis in the ESI mode was carried out on a Waters Acquity-Xevo G2-XS QTof.

### Preparation Procedure and Characterization Data for DSF and DS

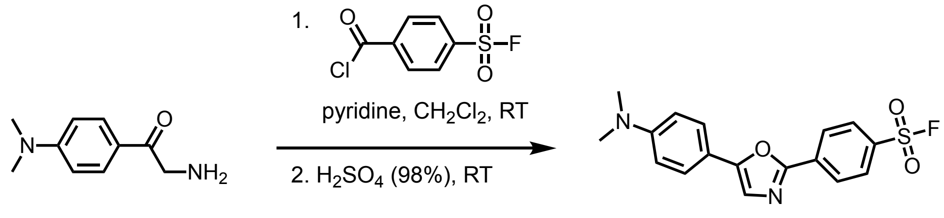

**Synthesis of 4-(5-(4-(dimethylamino)phenyl)oxazol-2-yl)benzenesulfonyl fluoride (dapoxyl sulfonyl fluoride, DSF):** In a 100 mL round bottomed flask, 2-amino-4'-dimethylaminoacetophenone (247 mg, 1.15 mmol, 1.15 equiv) and 4-(fluorosulfonyl)benzoyl chloride (1.00 mmol, 1.00 eq, 247 mg) were dissolved in 50 mL methylene chloride and anhydrous pyridine (3.00 mmol, 3.00 eq, 241 μL) was added dropwise. The orange heterogenous mixture was stirred overnight at room temperature (RT) and the solvent was evaporated under reduced pressure. The resulting orange powder was dissolved in 10 mL $H_2SO_4$ (98%) and stirred for two hours at RT. The clear orange solution was poured into a flask containing ice cubes and mixed, which was then neutralized with saturated $Na_2CO_3$ solution and then centrifuged. The crude product was collected by decantation and purified by silica gel column chromatography ($CH_2Cl_2$/MeOH step gradient). Concentration of the product fractions afforded DSF as a yellow solid (210 mg, 0.60 mmol, 60% yield, ≥95 % purity by NMR).

**$^1$H NMR** (500 MHz, $CDCl_3$) δ 8.29 (d, $J$ = 8.4 Hz, 2H), 8.09 (d, $J$ = 8.4 Hz, 2H), 7.61 (d, $J$ = 8.6 Hz, 2H), 7.34 (s, 1H), 6.77 (d, $J$ = 8.6, 2H), 3.04 (s, 6H).

**$^{13}$C NMR** (126 MHz, $CDCl_3$) δ 157.3, 154.1, 150.9, 134.1, 133.0, 132.8, 129.0, 126.5, 125.9, 121.6, 115.0, 112.1, 40.2.

**$^{19}$F NMR** (471 MHz, $CDCl_3$) δ 66.3.

**HRMS** (ESI) m/z: $[M + H]^+$ Calculated for $C_{17}H_{16}FN_2O_3S^+$, 347.0860; found 347.0864.

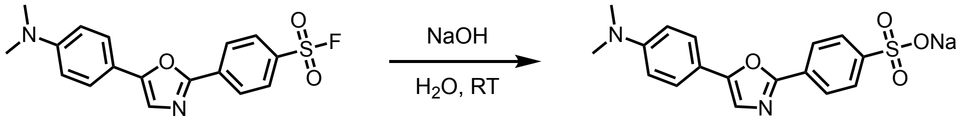

dapoxyl sulfonyl fluoride    sodium dapoxyl sulfonate (DS)

**Synthesis of sodium 4-(5-(4-(dimethylamino)phenyl)oxazol-2-yl)benzenesulfonate (sodium dapoxyl sulfonate, DS):** In a 25-mL round-bottom flask, DSF (15.0 mg, 0.0433 mmol) was suspended with 10% aqueous NaOH (5 mL) and refluxed (110 ºC) overnight while stirring. The mixture was then cooled down to RT and neutralized with 1 M HCl. The resulting yellow solid was chilled in an ice bath and recovered by decantation, followed by centrifugation. The isolated DS precipitate was dissolved in Milli-Q® water. The solution was filtered and lyophilized. The crude product was collected by filtration and purified by reversed-phase column chromatography (water / acetonitrile, step gradient from 0 to 100% acetonitrile). Fractions containing the product were lyophilized to afford DS as pale yellow powder (10.5 mg, 0.0287 mmol, 66% yield, ≥95 % purity by NMR).

**$^1$H NMR** (500 MHz, $d_6$-DMSO) δ 7.99 (d, *J* = 8.4 Hz 2H), 7.73 (d, *J* = 8.5 Hz, 2H), 7.65 (d, *J* = 8.9 Hz, 2H), 7.55 (s, 1H), 6.81 (d, *J* = 8.9, Hz, 2H), and 2.98 (s, 6H).

**$^{13}$C NMR** (126 MHz, $d_6$-DMSO) δ 158.6, 151.9, 150.4, 149.7, 126.9, 126.3, 125.3, 125.0, 121.0, 115.0, 112.2, and 39.8.

**HRMS** (ESI) m/z: $[M + H]^+$ Calculated for $C_{17}H_{17}N_2O_4S^+$ 345.0904; found 345.0841.

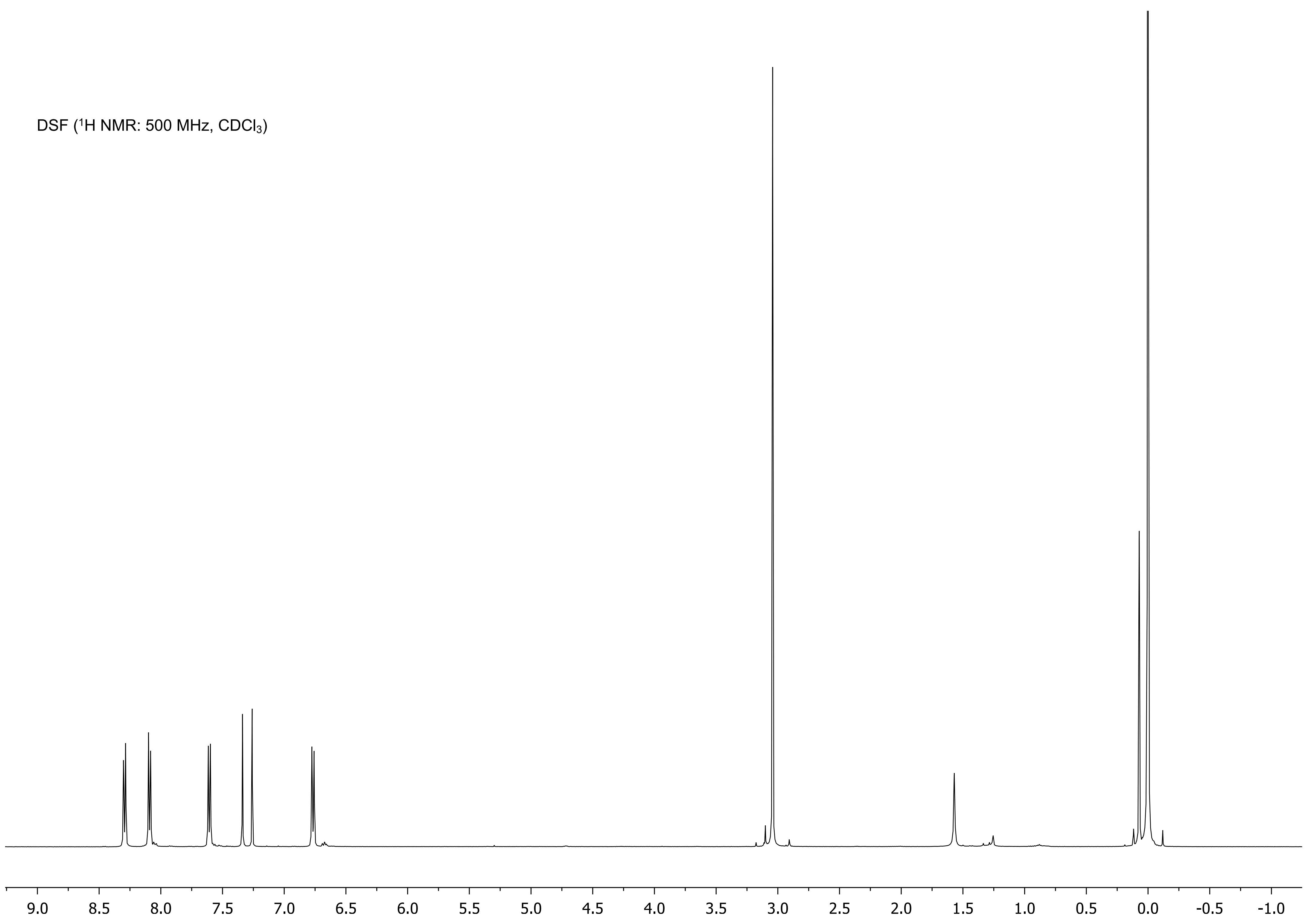
DSF ($^1$H NMR: 500 MHz, $CDCl_3$)
9.0 8.5 8.0 7.5 7.0 6.5 6.0 5.5 5.0 4.5 4.0 3.5 3.0 2.5 2.0 1.5 1.0 0.5 0.0 -0.5 -1.0

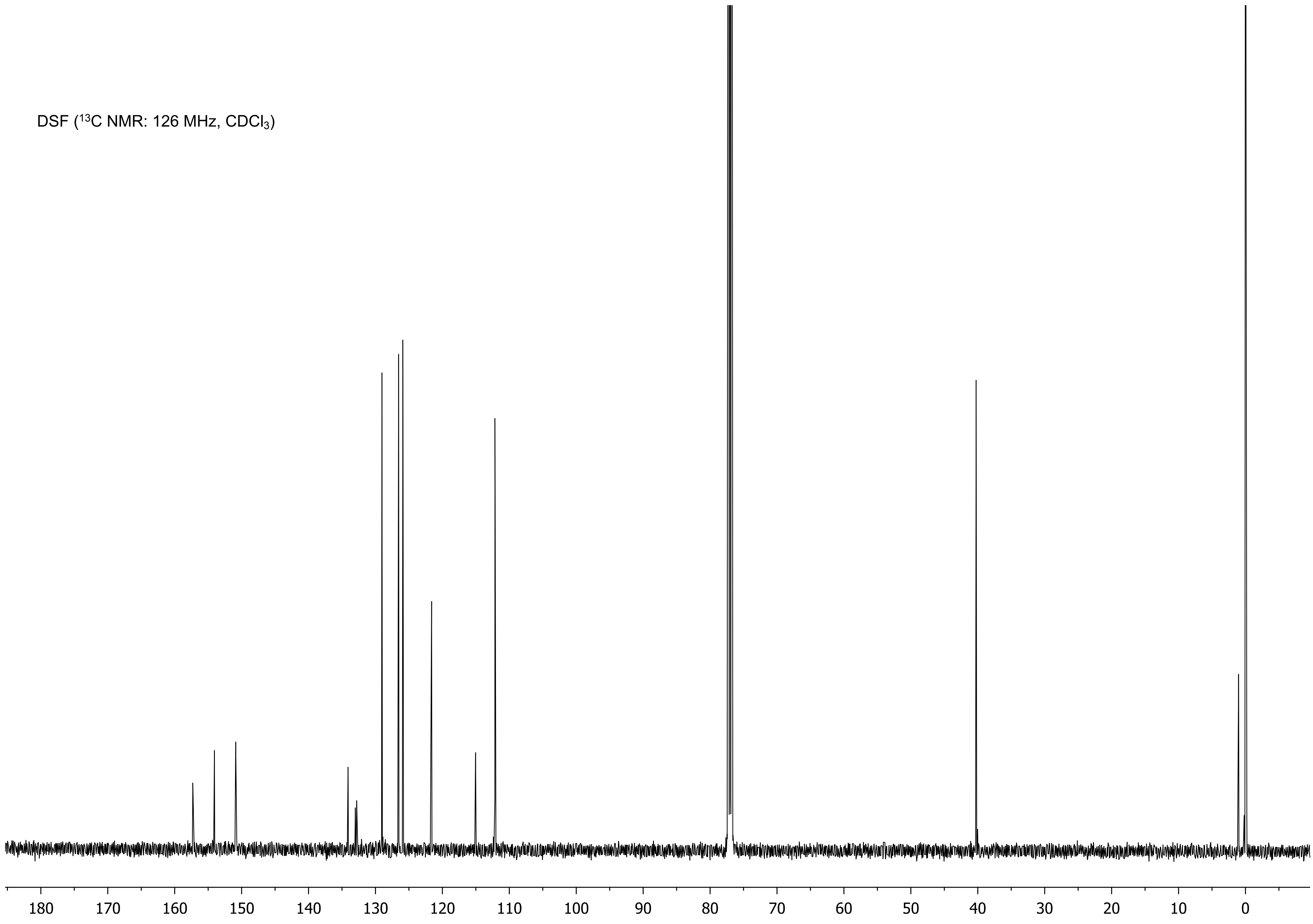
DSF ($^{13}C$ NMR: 126 MHz, $CDCl_3$)
180 170 160 150 140 130 120 110 100 90 80 70 60 50 40 30 20 10 0

DSF ($^{19}$F NMR: 471 MHz, $CDCl_3$)

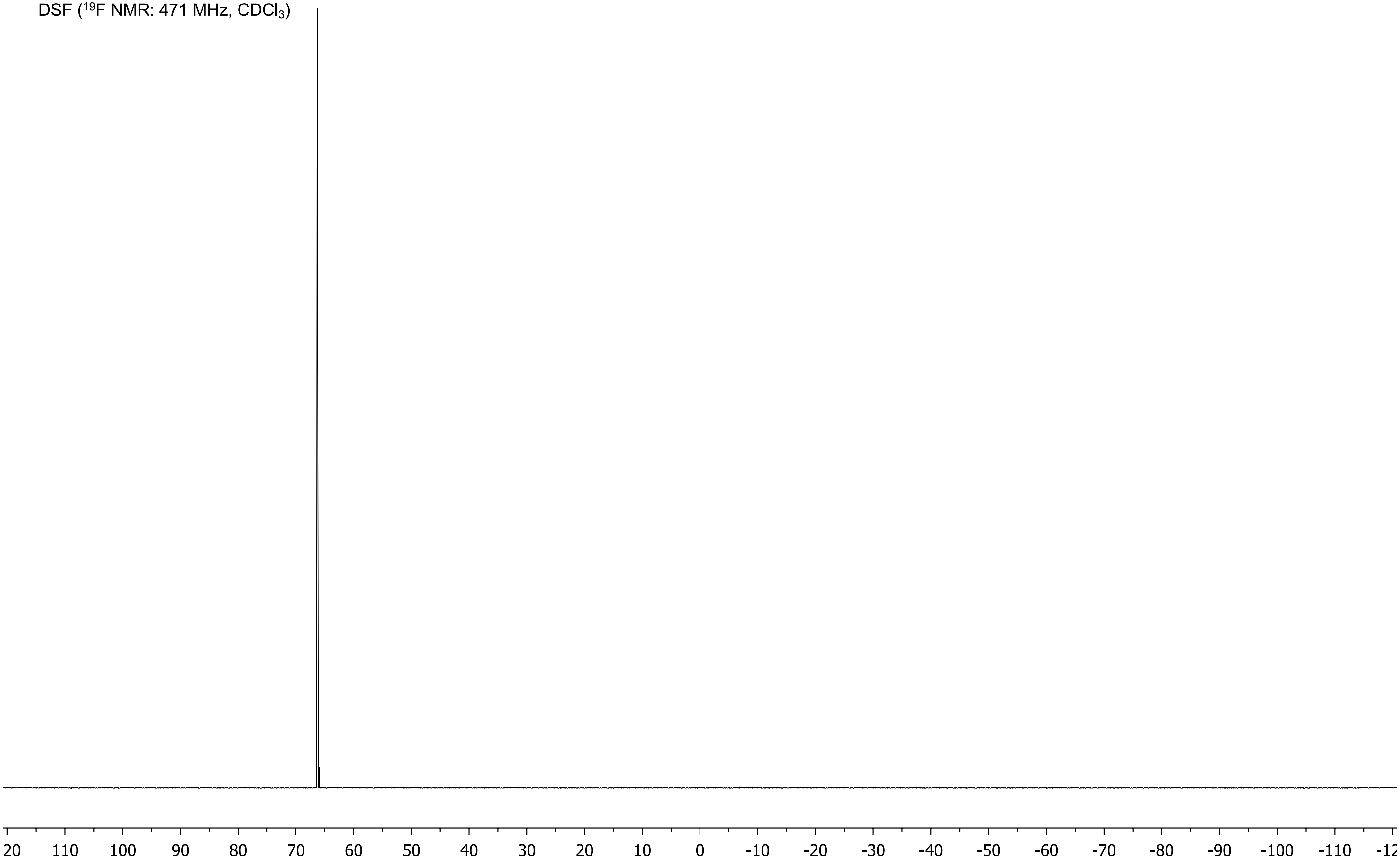

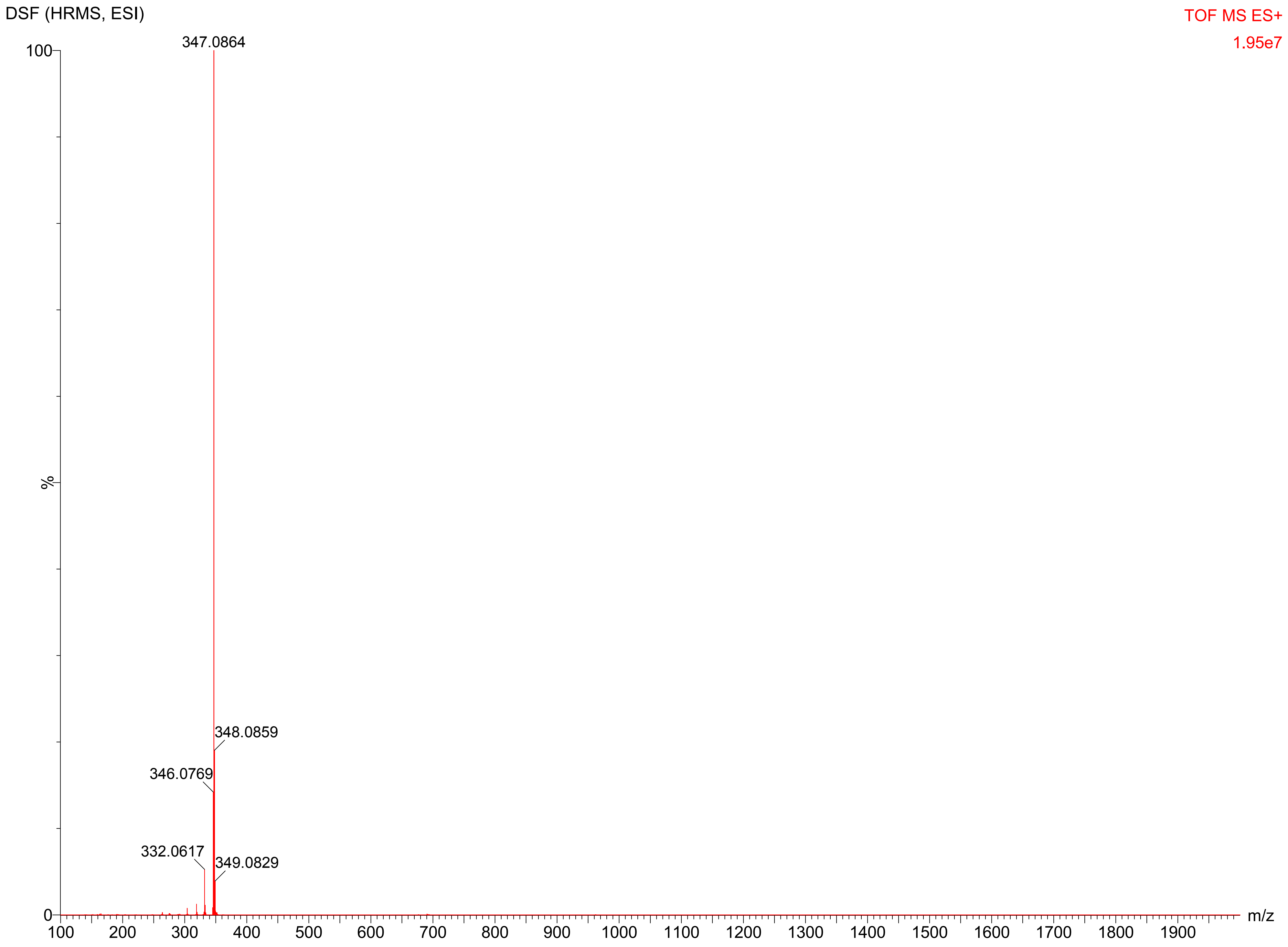

DSF (HRMS, ESI)
TOF MS ES+
1.95e7
347.0864
348.0859
346.0769
332.0617
349.0829
100
%
0
m/z
100
200
300
400
500
600
700
800
900
1000
1100
1200
1300
1400
1500
1600
1700
1800
1900

DS ($^1H$ NMR: 500 MHz, $d_6$-DMSO)

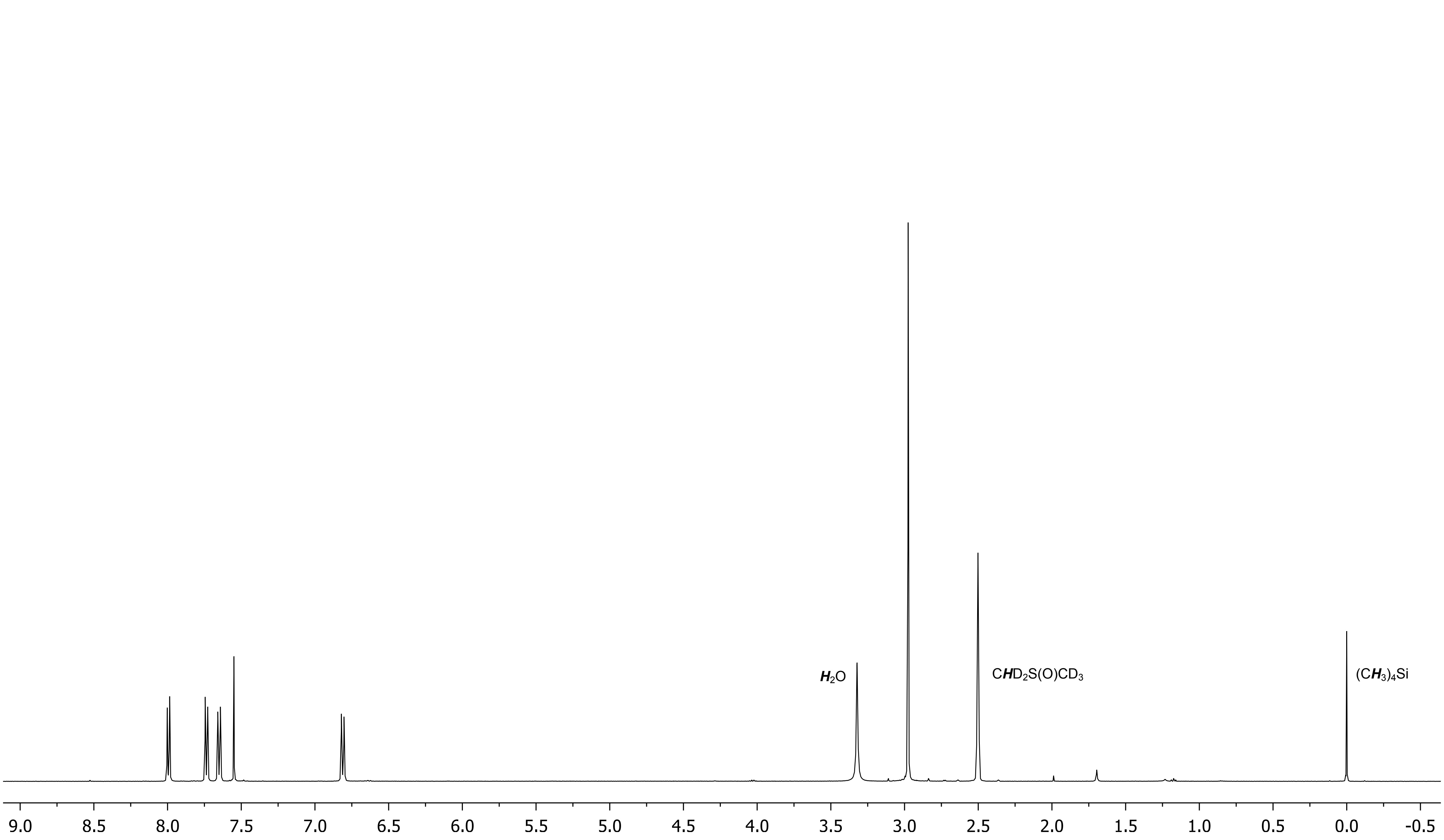

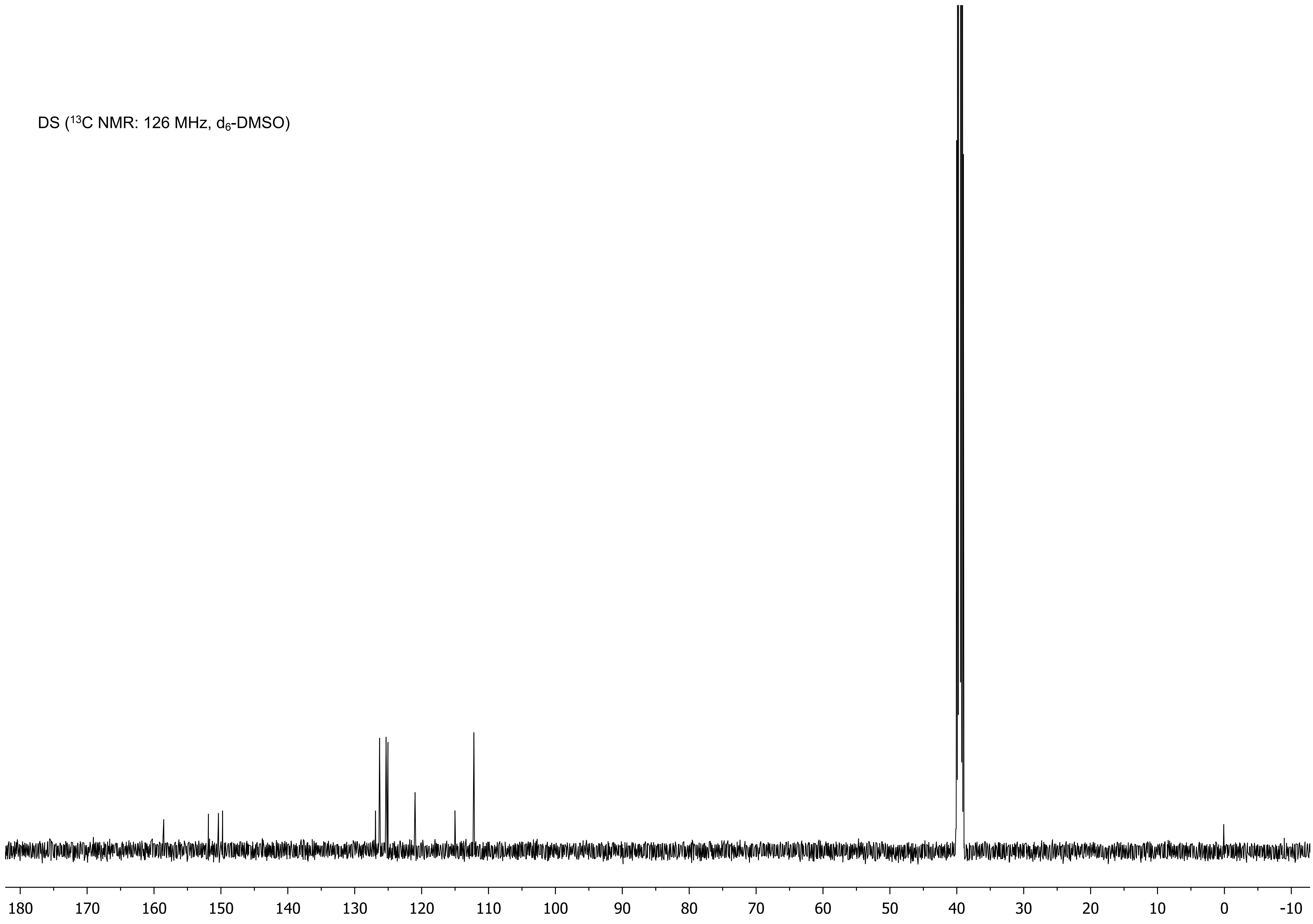

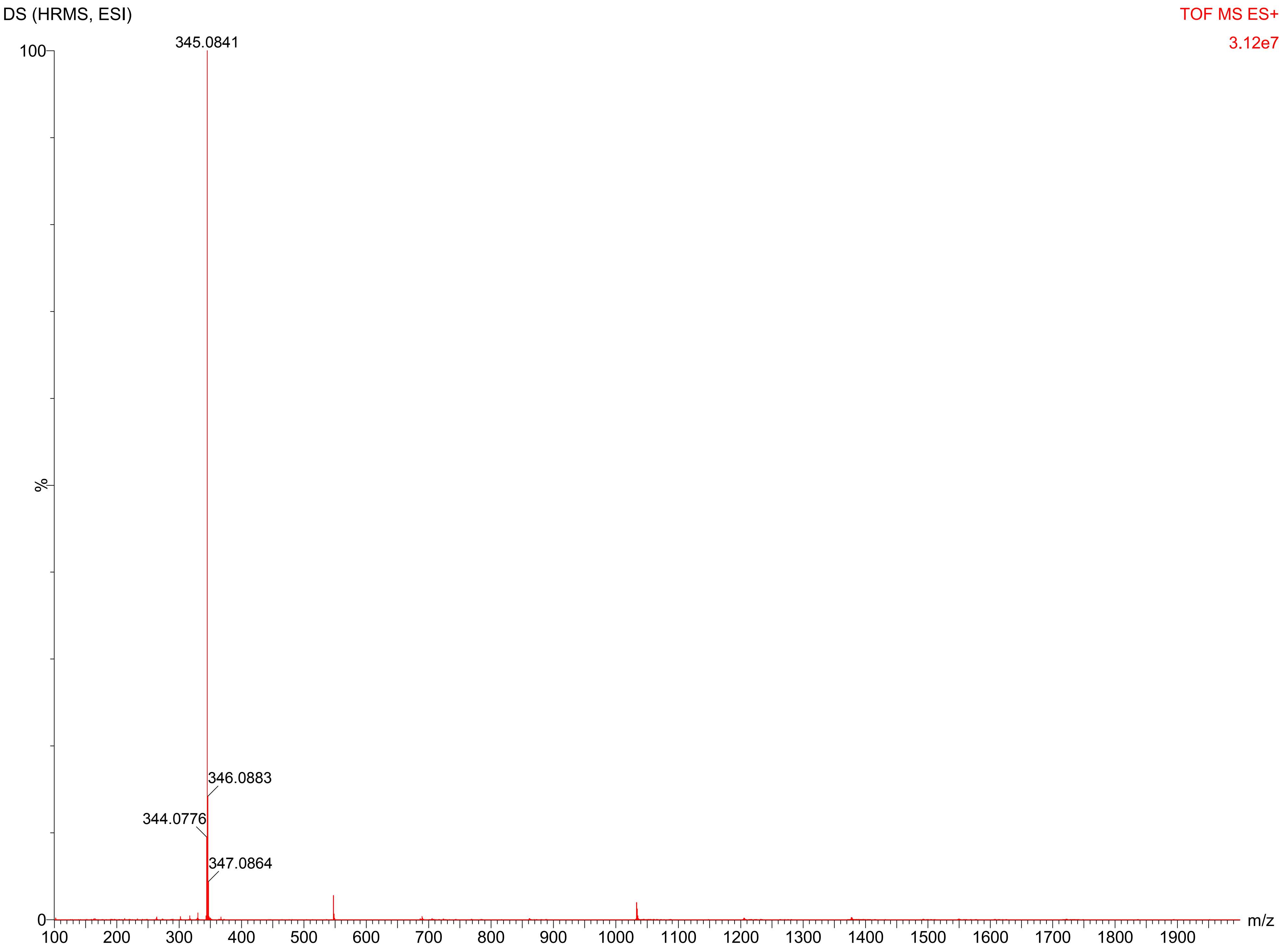
DS (HRMS, ESI)
TOF MS ES+
3.12e7
345.0841
346.0883
344.0776
347.0864
100
%
0
m/z
100
200
300
400
500
600
700
800
900
1000
1100
1200
1300
1400
1500
1600
1700
1800
1900

### Preparation and $^{1}$H NMR Quantification of AO

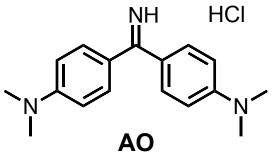

**AO**

To a 1.5 mL Eppendorf tube was placed AO (126 mg, 85% dye content per manufacturer) and ACS grade acetone (200 µL). The suspension was vortexed for 5 min and centrifuged to settle the solid. The supernatant was removed, and the solid was sequentially washed with acetone (200 µL) twice and diethyl ether (200 µL). The isolated solid was left under reduced pressure to remove residual organic volatiles, which provided a final amount of 83 mg of AO. For the purity assessment, 3.0 mg of AO and 7.6 mg (62.3 mmol) of benzoic acid (≥99% per manufacturer) as an internal standard were dissolved in 350 µL of $d_6$-DMSO. The purity of AO was determined to be ≥98% by $^{1}$H NMR spectroscopy (see below). A 5 mM stock solution of AO was then made by dissolving 10 mg in 6.6 mL of Milli-Q® water. Depending on the fluorescence assay, this stock solution was diluted with Milli-Q® water to prepare a secondary stock solution.

AO ([1]H NMR: 500 MHz, $d_6$-DMSO)

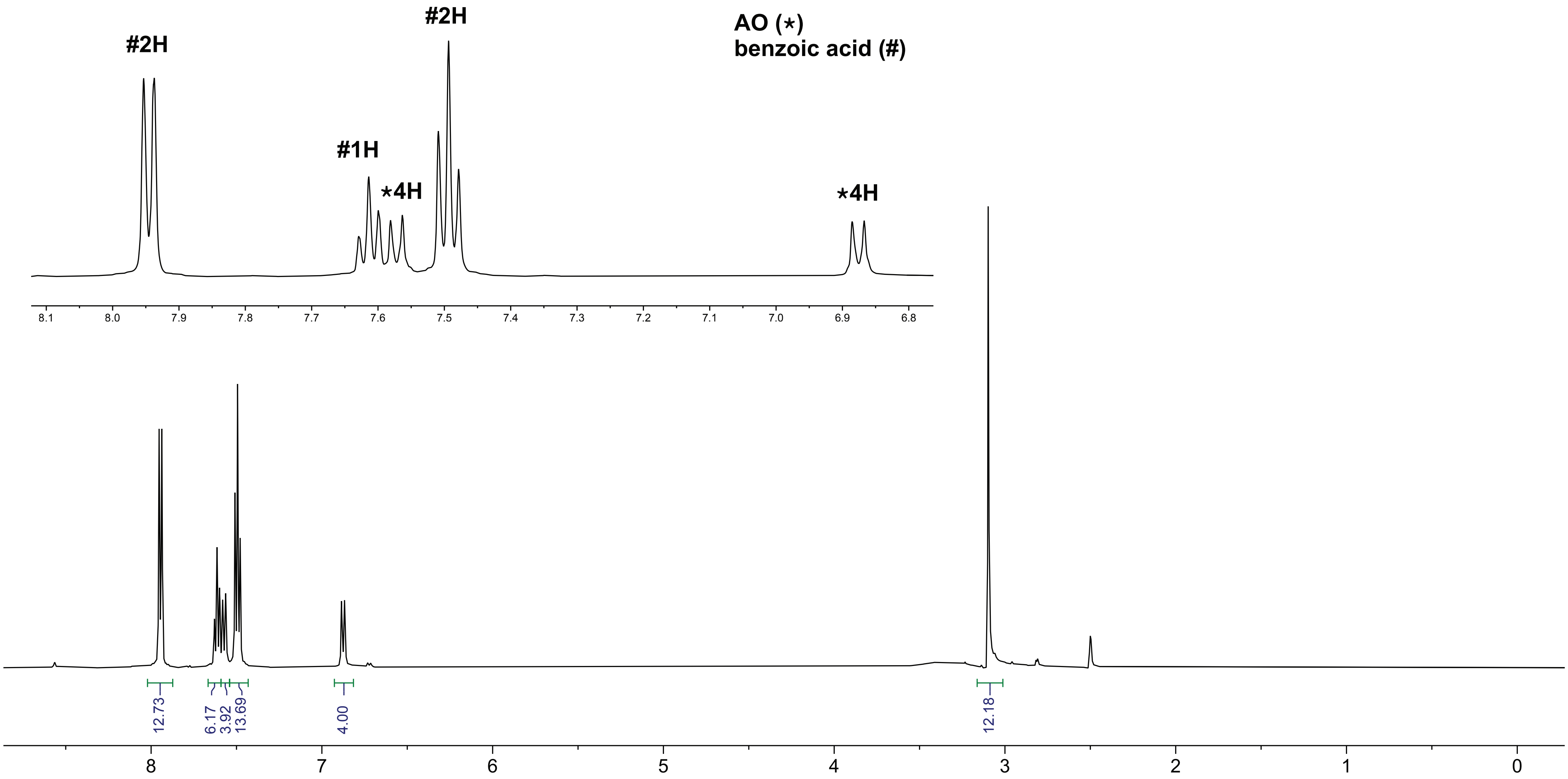

### MiRNA Secondary Structures and Hybridization Energies

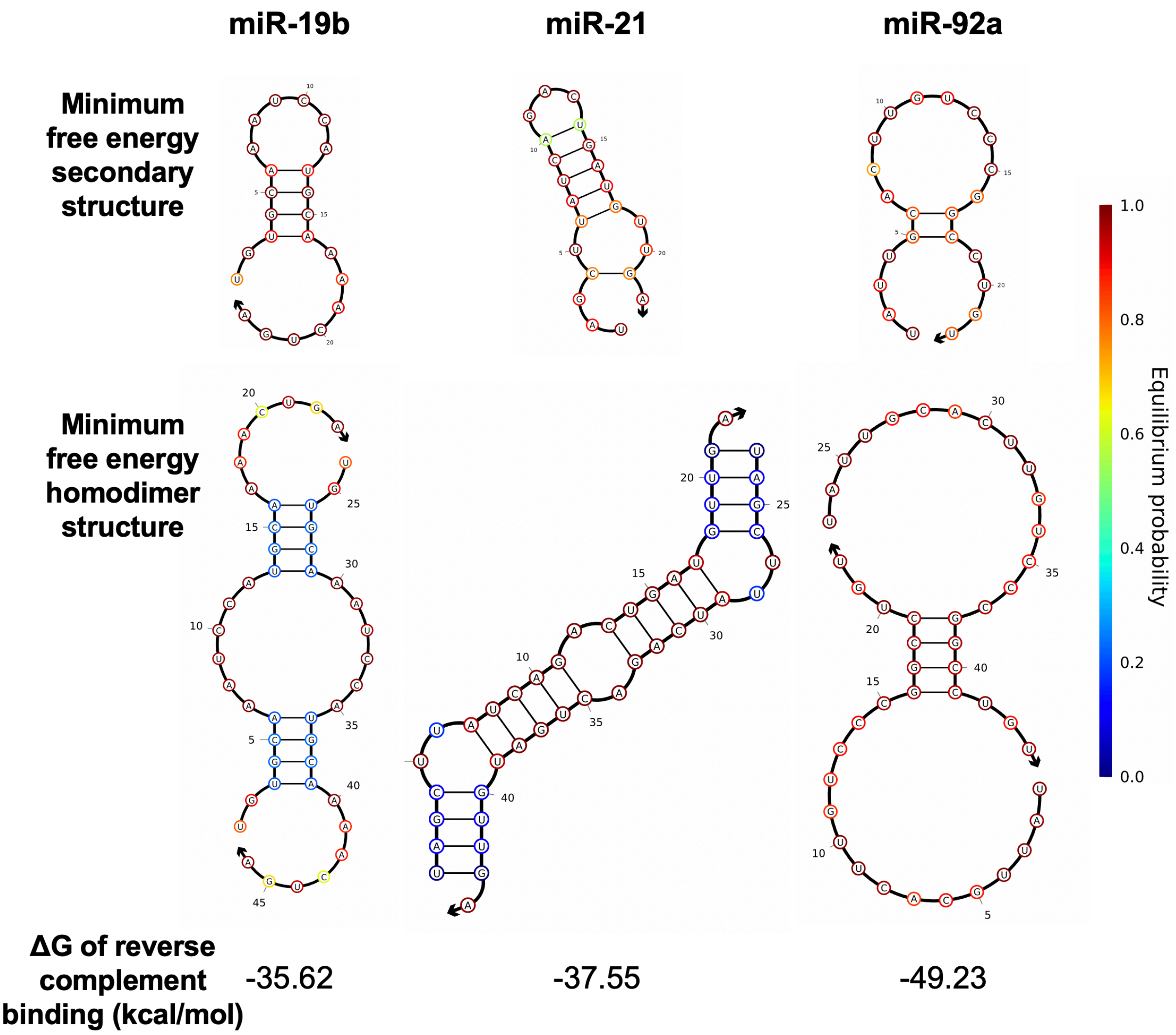

**Supplementary Fig. 1.** Comparison of secondary structures and hybridization energies of miRNAs used in this study. MiR-19b exhibits the strongest secondary structure. miR-21 forms a homodimer with high equilibrium probability. miR-19 has the highest binding energy to a complementary sequence. Visualization was performed with NUPACK[2]. ΔG calculations were performed with Vienna RNA Websuite[3].

**Comparative Analysis of Fluorogenicity of DSF, DS, and AO in Buffer and Biological Media**

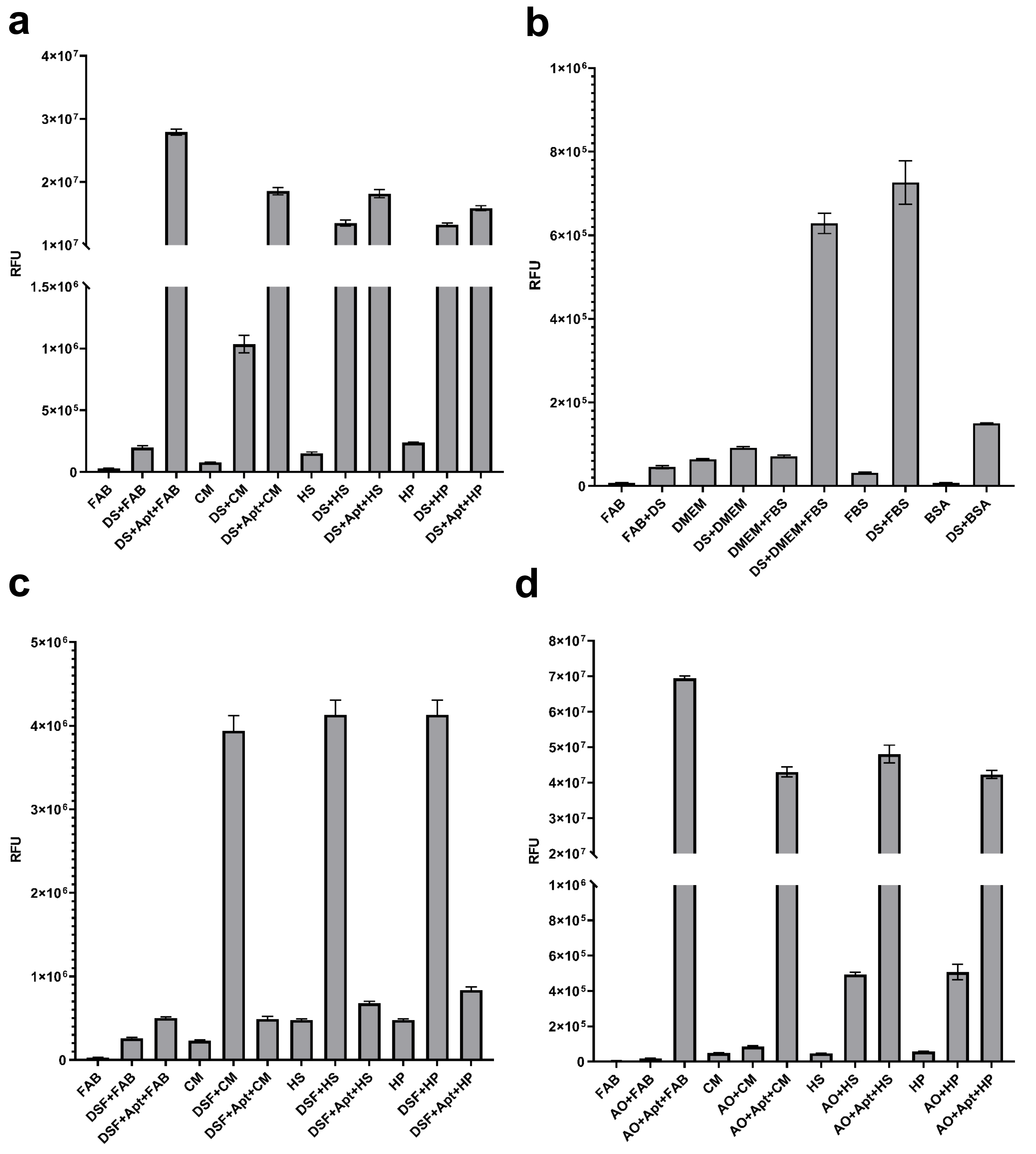

**Supplementary Fig. 2.** Comparison of fluorescence intensity of (**a**-**b**) DS, (**c**) DSF, and (**d**) AO in fluorescence assay buffer (FAB) and biological media (10%). 10 μM fluorophore, 2 μM DAP-10 (5′-CAATTACGGGGGAGGGTGTGTGGTCTTGCTTGGTTCGTATTG), and 10% (v/v) media were used. Fluorescence intensities were measured at the following $\lambda_{ex}$ / $\lambda_{em}$: DS: 391 / 497 nm; DSF: 390 / 505 nm; AO: 475 / 540 nm. Error bars denote the standard deviation of three replicates. Final concentrations of DMEM: 10%, FBS: 1% v/v, BSA: 0.025 mg/mL.

### MiRNAs and MiRNA Mutants

The complete list of oncogenic miRNAs and mutant miRNAs used in this study is provided in Supplementary Table 1. Mutants include single nucleotide variation compared to the native miRNA and are classified as *NXM*, where *N* designates the native nucleotide, *X* is the nucleotide number in the sequence, and *M* designates the nucleotide variant.

**Supplementary Table 1.** Sequences of miRNAs and miRNA mutants.

| miRNA | Mutation | Sequence (5′ to 3′) |
|---|---|---|
| miR-19b | None | UGUGCAAAUCCAUGCAAAACUGA |
| | G2A | UAUGCAAAUCCAUGCAAAACUGA |
| | U3G | UGGGCAAAUCCAUGCAAAACUGA |
| | G4A | UGUACAAAUCCAUGCAAAACUGA |
| | C5G | UGUGGAAAUCCAUGCAAAACUGA |
| | A6U | UGUGCUAAUCCAUGCAAAACUGA |
| | A16U | UGUGCAAAUCCAUGCUAAACUGA |
| | A17U | UGUGCAAAUCCAUGCAUAACUGA |
| | A18U | UGUGCAAAUCCAUGCAAUACUGA |
| | A19U | UGUGCAAAUCCAUGCAAAUCUGA |
| | C20G | UGUGCAAAUCCAUGCAAAAGUGA |
| | U21G | UGUGCAAAUCCAUGCAAAACGGA |
| miR-21 | None | UAGCUUAUCAGACUGAUGUUGA |
| | A2U | UUGCUUAUCAGACUGAUGUUGA |
| | G3A | UAACUUAUCAGACUGAUGUUGA |
| | C4G | UAGGUUAUCAGACUGAUGUUGA |
| | U5G | UAGCGUAUCAGACUGAUGUUGA |
| | U6G | UAGCUGAUCAGACUGAUGUUGA |
| | A16U | UAGCUUAUCAGACUGUUGUUGA |
| | U17G | UAGCUUAUCAGACUGAGGUUGA |
| | G18A | UAGCUUAUCAGACUGAUAUUGA |
| | U19G | UAGCUUAUCAGACUGAUGGUGA |
| | U20G | UAGCUUAUCAGACUGAUGUGGA |
| | G21A | UAGCUUAUCAGACUGAUGUUAA |
| miR-92a | None | UAUUGCACUUGUCCCGGCCUGU |
| | A2U | UUUUGCACUUGUCCCGGCCUGU |
| | U3G | UAGUGCACUUGUCCCGGCCUGU |
| | U4G | UAUGGCACUUGUCCCGGCCUGU |
| | G5A | UAUUACACUUGUCCCGGCCUGU |

| | | |
|---|---|---|
| | C6G | UAUUGGACUUGUCCCGGCCUGU |
| | G16A | UAUUGCACUUGUCCCAGCCUGU |
| | G17A | UAUUGCACUUGUCCCGACCUGU |
| | C18G | UAUUGCACUUGUCCCGGGCUGU |
| | C19G | UAUUGCACUUGUCCCGGCGUGU |
| | U20G | UAUUGCACUUGUCCCGGCCGGU |
| | G21A | UAUUGCACUUGUCCCGGCCUAU |

### Split Aptamer Design

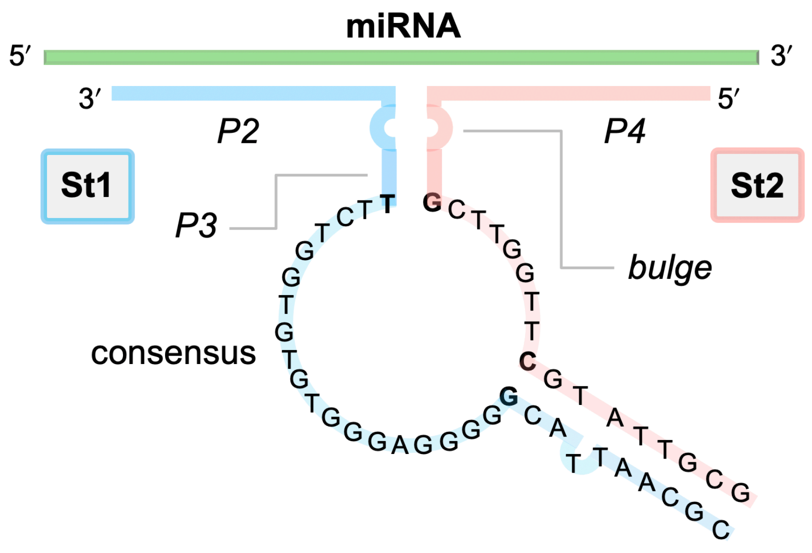

**Supplementary Fig. 3.** Design of **St1:St2:miRNA**.

**Supplementary Table 2.** St1:St2 Designations.

| Letter Designation | P2 (nts) | P4 (nts) | 35G |
|---|---|---|---|
| a | 9 | 10 | + |
| b | 9 | 9 | + |
| c | 9 | 9 | - |
| d | 8 | 9 | - |
| e | 8 | 10 | - |

| Numeric Designation | bulge (nts) | P3 (nts) |
|---|---|---|
| 1 | 2 | 3 |
| 2 | 1 | 3 |
| 3 | 0 | 3 |
| 4 | 2 | 2 |
| 5 | 1 | 2 |
| 6 | 2 | 1 |
| 7 | 1 | 1 |
| 8 | 0 | 0 |

**Supplementary Table 3.** DNA aptamer precursor strands **St1** and **St2**.

| ID | Aptamer DNA Precursor Sequence (5′ to 3′) St1 | St2 |
|---|---|---|
| ***Targeting miR-19b*** | | |
| a1 | CGCAATTACGGGGGAGGGTGTGTGGTCTTGTCTTGGATTTGCA | AGTTTTGCATTTGACGCTTGGTTCGTATTGCG |
| a2 | CGCAATTACGGGGGAGGGTGTGTGGTCTTGTCTGGATTTGCA | AGTTTTGCATTGACGCTTGGTTCGTATTGCG |
| a3 | CGCAATTACGGGGGAGGGTGTGTGGTCTTGTCGGATTTGCA | AGTTTTGCATGACGCTTGGTTCGTATTGCG |
| a4 | CGCAATTACGGGGGAGGGTGTGTGGTCTTGTTTGGATTTGCA | AGTTTTGCATTTACGCTTGGTTCGTATTGCG |
| a5 | CGCAATTACGGGGGAGGGTGTGTGGTCTTGTTGGATTTGCA | AGTTTTGCATTACGCTTGGTTCGTATTGCG |
| a6 | CGCAATTACGGGGGAGGGTGTGTGGTCTTGTTGGATTTGCA | AGTTTTGCATTTCGCTTGGTTCGTATTGCG |
| a7 | CGCAATTACGGGGGAGGGTGTGTGGTCTTGTGGATTTGCA | AGTTTTGCATTCGCTTGGTTCGTATTGCG |
| a8 | CGCAATTACGGGGGAGGGTGTGTGGTCTTGGATTTGCA | AGTTTTGCATGCTTGGTTCGTATTGCG |
| b1 | CGCAATTACGGGGGAGGGTGTGTGGTCTTGTCTTGGATTTGCA | GTTTTGCATTTGACGCTTGGTTCGTATTGCG |
| b2 | CGCAATTACGGGGGAGGGTGTGTGGTCTTGTCTGGATTTGCA | GTTTTGCATTTGACGCTTGGTTCGTATTGCG |
| b3 | CGCAATTACGGGGGAGGGTGTGTGGTCTTGTCGGATTTGCA | GTTTTGCATTGACGCTTGGTTCGTATTGCG |
| b4 | CGCAATTACGGGGGAGGGTGTGTGGTCTTGTTTGGATTTGCA | GTTTTGCATTTACGCTTGGTTCGTATTGCG |
| b5 | CGCAATTACGGGGGAGGGTGTGTGGTCTTGTTGGATTTGCA | GTTTTGCATTACGCTTGGTTCGTATTGCG |
| b6 | CGCAATTACGGGGGAGGGTGTGTGGTCTTGTTGGATTTGCA | GTTTTGCATTTCGCTTGGTTCGTATTGCG |
| b7 | CGCAATTACGGGGGAGGGTGTGTGGTCTTGTGGATTTGCA | GTTTTGCATTCGCTTGGTTCGTATTGCG |
| b8 | CGCAATTACGGGGGAGGGTGTGTGGTCTTGGATTTGCA | GTTTTGCATGCTTGGTTCGTATTGCG |
| c1 | CGCAATTACGGGGGAGGGTGTGTGGTCTTGTCTTGATTTGCAC | GTTTTGCATTTGACGCTTGGTTCGTATTGCG |
| c2 | CGCAATTACGGGGGAGGGTGTGTGGTCTTGTCTGATTTGCAC | GTTTTGCATTTGACGCTTGGTTCGTATTGCG |
| c3 | CGCAATTACGGGGGAGGGTGTGTGGTCTTGTCGATTTGCAC | GTTTTGCATTGACGCTTGGTTCGTATTGCG |
| c4 | CGCAATTACGGGGGAGGGTGTGTGGTCTTGTTTGATTTGCAC | GTTTTGCATTTACGCTTGGTTCGTATTGCG |
| c5 | CGCAATTACGGGGGAGGGTGTGTGGTCTTGTTGATTTGCAC | GTTTTGCATTACGCTTGGTTCGTATTGCG |
| c6 | CGCAATTACGGGGGAGGGTGTGTGGTCTTGTTGATTTGCAC | GTTTTGCATTTCGCTTGGTTCGTATTGCG |
| c7 | CGCAATTACGGGGGAGGGTGTGTGGTCTTGTGATTTGCAC | GTTTTGCATTCGCTTGGTTCGTATTGCG |
| c8 | CGCAATTACGGGGGAGGGTGTGTGGTCTTGATTTGCAC | GTTTTGCATGCTTGGTTCGTATTGCG |
| d1 | CGCAATTACGGGGGAGGGTGTGTGGTCTTGTCTTGATTTGCA | GTTTTGCATTTGACGCTTGGTTCGTATTGCG |
| d2 | CGCAATTACGGGGGAGGGTGTGTGGTCTTGTCTGATTTGCA | GTTTTGCATTTGACGCTTGGTTCGTATTGCG |
| d3 | CGCAATTACGGGGGAGGGTGTGTGGTCTTGTCGATTTGCA | GTTTTGCATTGACGCTTGGTTCGTATTGCG |
| d4 | CGCAATTACGGGGGAGGGTGTGTGGTCTTGTTTGATTTGCA | GTTTTGCATTTACGCTTGGTTCGTATTGCG |
| d5 | CGCAATTACGGGGGAGGGTGTGTGGTCTTGTTGATTTGCA | GTTTTGCATTACGCTTGGTTCGTATTGCG |
| d6 | CGCAATTACGGGGGAGGGTGTGTGGTCTTGTTGATTTGCA | GTTTTGCATTTCGCTTGGTTCGTATTGCG |
| d7 | CGCAATTACGGGGGAGGGTGTGTGGTCTTGTGATTTGCA | GTTTTGCATTCGCTTGGTTCGTATTGCG |
| d8 | CGCAATTACGGGGGAGGGTGTGTGGTCTTGATTTGCA | GTTTTGCATGCTTGGTTCGTATTGCG |
| e1 | CGCAATTACGGGGGAGGGTGTGTGGTCTTGTCTTGATTTGCA | AGTTTTGCATTTGACGCTTGGTTCGTATTGCG |
| e2 | CGCAATTACGGGGGAGGGTGTGTGGTCTTGTCTGATTTGCA | AGTTTTGCATTGACGCTTGGTTCGTATTGCG |
| e3 | CGCAATTACGGGGGAGGGTGTGTGGTCTTGTCGATTTGCA | AGTTTTGCATGACGCTTGGTTCGTATTGCG |
| e4 | CGCAATTACGGGGGAGGGTGTGTGGTCTTGTTTGATTTGCA | AGTTTTGCATTTACGCTTGGTTCGTATTGCG |
| e5 | CGCAATTACGGGGGAGGGTGTGTGGTCTTGTTGATTTGCA | AGTTTTGCATTACGCTTGGTTCGTATTGCG |

| | | |
|---|---|---|
| e6 | CGCAATTACGGGGGAGGGTGTGTGGTCTTGTTGATTTGCA | AGTTTTGCATTTCGCTTGGTTCGTATTGCG |
| e7 | CGCAATTACGGGGGAGGGTGTGTGGTCTTGTGATTTGCA | AGTTTTGCATTCGCTTGGTTCGTATTGCG |
| e8 | CGCAATTACGGGGGAGGGTGTGTGGTCTTGATTTGCA | AGTTTTGCATGCTTGGTTCGTATTGCG |
| ***Targeting miR-21*** | | |
| a1 | CGCAATTACGGGGGAGGGTGTGTGGTCTTGTCTTCTGATAAGC | CAACATCAGTTTGACGCTTGGTTCGTATTGCG |
| a2 | CGCAATTACGGGGGAGGGTGTGTGGTCTTGTCTCTGATAAGC | CAACATCAGTTGACGCTTGGTTCGTATTGCG |
| a6 | CGCAATTACGGGGGAGGGTGTGTGGTCTTGTTCTGATAAGC | CAACATCAGTTTCGCTTGGTTCGTATTGCG |
| a7 | CGCAATTACGGGGGAGGGTGTGTGGTCTTGTCTGATAAGC | CAACATCAGTTCGCTTGGTTCGTATTGCG |
| a8 | CGCAATTACGGGGGAGGGTGTGTGGTCTTCTGATAAGC | CAACATCAGTGCTTGGTTCGTATTGCG |
| b1 | CGCAATTACGGGGGAGGGTGTGTGGTCTTGTCTTCTGATAAGC | AACATCAGTTTGACGCTTGGTTCGTATTGCG |
| b7 | CGCAATTACGGGGGAGGGTGTGTGGTCTTGTCTGATAAGC | AACATCAGTTCGCTTGGTTCGTATTGCG |
| b8 | CGCAATTACGGGGGAGGGTGTGTGGTCTTCTGATAAGC | AACATCAGTGCTTGGTTCGTATTGCG |
| c1 | CGCAATTACGGGGGAGGGTGTGTGGTCTTGTCTTTGATAAGCT | AACATCAGTTTGACGCTTGGTTCGTATTGCG |
| c2 | CGCAATTACGGGGGAGGGTGTGTGGTCTTGTCTTGATAAGCT | AACATCAGTTGACGCTTGGTTCGTATTGCG |
| c3 | CGCAATTACGGGGGAGGGTGTGTGGTCTTGTCTGATAAGCT | AACATCAGTGACGCTTGGTTCGTATTGCG |
| c4 | CGCAATTACGGGGGAGGGTGTGTGGTCTTGTTTTGATAAGCT | AACATCAGTTTACGCTTGGTTCGTATTGCG |
| ***Targeting miR-92a*** | | |
| a1 | CGCAATTACGGGGGAGGGTGTGTGGTCTTGTCTTCAAGTGCAA | CAGGCCGGGATTGACGCTTGGTTCGTATTGCG |
| a2 | CGCAATTACGGGGGAGGGTGTGTGGTCTTGTCTCAAGTGCAA | CAGGCCGGGATGACGCTTGGTTCGTATTGCG |
| a6 | CGCAATTACGGGGGAGGGTGTGTGGTCTTGTTCAAGTGCAA | CAGGCCGGGATTCGCTTGGTTCGTATTGCG |
| a7 | CGCAATTACGGGGGAGGGTGTGTGGTCTTGTCAAGTGCAA | CAGGCCGGGATCGCTTGGTTCGTATTGCG |
| a8 | CGCAATTACGGGGGAGGGTGTGTGGTCTTCAAGTGCAA | CAGGCCGGGAGCTTGGTTCGTATTGCG |
| b1 | CGCAATTACGGGGGAGGGTGTGTGGTCTTGTCTTCAAGTGCAA | AGGCCGGGATTGACGCTTGGTTCGTATTGCG |
| b7 | CGCAATTACGGGGGAGGGTGTGTGGTCTTGTCAAGTGCAA | AGGCCGGGATCGCTTGGTTCGTATTGCG |
| b8 | CGCAATTACGGGGGAGGGTGTGTGGTCTTCAAGTGCAA | AGGCCGGGAGCTTGGTTCGTATTGCG |
| c1 | CGCAATTACGGGGGAGGGTGTGTGGTCTTGTCTTAAGTGCAAT | AGGCCGGGATTGACGCTTGGTTCGTATTGCG |
| c2 | CGCAATTACGGGGGAGGGTGTGTGGTCTTGTCTAAGTGCAAT | AGGCCGGGATGACGCTTGGTTCGTATTGCG |
| c3 | CGCAATTACGGGGGAGGGTGTGTGGTCTTGTCAAGTGCAAT | AGGCCGGGAGACGCTTGGTTCGTATTGCG |
| c4 | CGCAATTACGGGGGAGGGTGTGTGGTCTTGTTTAAGTGCAAT | AGGCCGGGATTACGCTTGGTTCGTATTGCG |

### MiRNA Detection

DNA strands St1 and St2 (100 μM stock solutions) were mixed and diluted with FAB (volumes vary depending on the assay). The mixtures were incubated at 80 ºC for 5 minutes and then cooled to RT. The resulting solution was aliquoted into 96 well plates (Greiner Bio-one). To these wells were sequentially added stock solutions of AO (25 μM) and miRNA (20 μM). The final concentrations were the following: 2 μM St1, 2 μM St2, 10 μM AO, and 2 μM miRNA. Fluorescence intensity of the samples were measured at 475 nm excitation over one hour on a microplate reader (SpectraMax iD3, Molecular Devices, LLC). For the samples containing biological media, undiluted CM, HS, or HP was first incubated with murine RNase inhibitor at RT for 5 minutes. The resulting solution was then added into 96 well plates. The pre-incubated St1 and St2, followed by AO and miRNA solutions were added into these wells. The final concentrations were the following: 10% or 30% (v/v) CM, HS, or HP, 2 μM St1, 2 μM St2, 10 μM AO, 2 μM miRNA, and 1 unit/μL murine RNase inhibitor. Of note, fluorescence intensities were observed to typically decrease by ~30% in FAB upon addition of murine RNase inhibitor.

### Dissociation Constant Determination

The dissociation constant of AO binding to the LigBR, $K_d$, was calculated in the presence and absence of the target miRNA using fluorescence assays. To calculate the $K_d$ with miRNA, $K_{d,1}$, a mixture of St1, St2, and miRNA was incubated at 80 ºC for 5 minutes and then cooled to RT to form the aptamer. Next, the aptamer was diluted to concentrations ranging from 0.25 to 7 µM. To these samples was added AO (1 µM final concentration) at RT and fluorescence intensities were continuously recorded by the plate reader. The intensities were used to calculate $K_{d,1}$ by fitting by nonlinear regression analysis to the equation below, which was derived from 1-to-1 binding.

$$[Bound\ AO]_1 = \frac{F_1 - F_0}{F_{max}}[AO]_T = \frac{[Apt]_T + [AO]_T + K_{d,1}}{2} - \sqrt{\left(\frac{[Apt]_T + [AO]_T + K_{d,1}}{2}\right)^2 - [Apt]_T[AO]_T} \quad [1]$$

In this equation, $F_1$ represents the observed fluorescence intensity, $F_0$ is the fluorescence intensity of 1 µM AO, $F_{max}$ is the theoretical maximum fluorescence intensity based on a defined AO concentration, and $[Apt]_T$ and $[AO]_T$ are the total concentrations of the aptamer and AO, respectively. $[Bound\ AO]_1$ is the concentration of bound state of AO in the presence of miRNA.

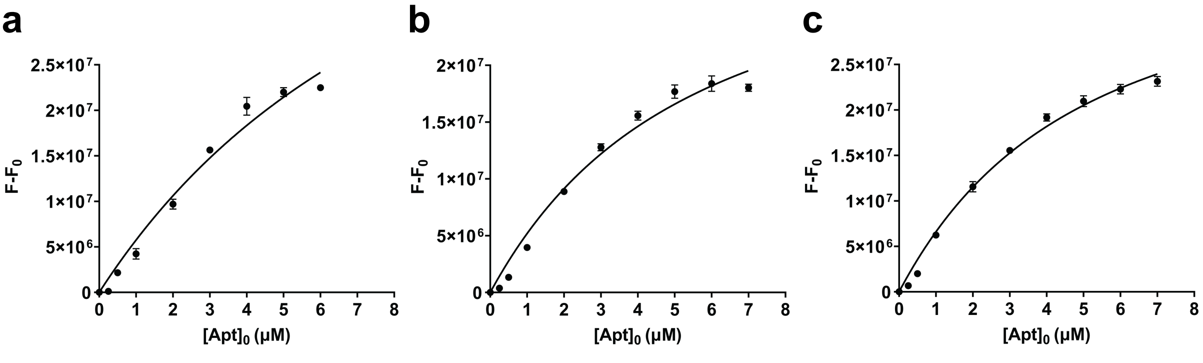

**Supplementary Fig. 4.** Determination of $K_{d,1}$ for complexes of AO and **(a)** c2-19b:miR-19b, **(b)** a2-21:miR-21, and **(c)** b7-92a:miR-92a. Error bars denote the standard deviation of three replicates.

Fitting by non-linear regression could not be used to determine $K_{d,2}$, which is the dissociation constant of AO binding to LigBR in the absence of the miRNA, because generation of adequate signal for curve fitting would require orders of magnitude higher St1 and St2 concentrations. To estimate the $K_{d,2}$, we used the $F_1/F_2$ obtained in the miRNA detection section. We first made 2 assumptions: 1) the fluorescence output of LigBR-bound AO is the same both in the presence and absence of the miRNA, and 2) the ratio of fluorescence intensity in the presence of miRNA to in the absence of miRNA is close to the ratio of bound AO in the presence of miRNA to in the absence of miRNA.

$$\frac{[Bound\ AO]_1}{[Bound\ AO]_2} \approx \frac{F_1}{F_2} \qquad [2]$$

In this equation, $[\text{Bound AO}]_2$ is the concentration of bound state of AO in the absence miRNA. Substituting equation [2] into equation [1], $K_{d,2}$ can be solved using the previously determined $K_{d,1}$.

$$[Bound\ AO]_1 \times \frac{F_2}{F_1} = \frac{[Apt]_T + [AO]_T + K_{d,2}}{2} - \sqrt{\left(\frac{[Apt]_T + [AO]_T + K_{d,2}}{2}\right)^2 - [Apt]_T[AO]_T}$$

**Supplementary Table 4.** Summary of data fitting.

| complex | $K_{d,1}$ (μM) | $R^2$ | St1:St2 | $F/F_0$ | $K_{d,2}$ (μM)* |
|---|---|---|---|---|---|
| c2-19b:miR-19b | 9.1 ± 2.8 | 0.9876 | c2-19b | 9.35 | 180 |
| a2-21:miR-21 | 4.6 ± 0.9 | 0.9804 | a2-21 | 9.84 | 140 |
| b7-92a:miR-92a | 4.2 ± 0.5 | 0.9912 | b7-92a | 21.36 | 310 |

* Estimated for AO binding to St1:St2 by employing $F/F_0$ and measured $K_d$.

### Sequence Specificity in Biological Media

Sequence specificities of the selected St1:St2 sets were assessed in the presence of 2 μM single nucleotide mutants that led to the lowest and highest differentiation factor, $D_f$, in FAB. Larger $D_f$ indicates better specificity. These mutants were then tested in biological media and the results are presented in bar graphs below. Samples prepared with biological media contained murine RNase inhibitor as described above.

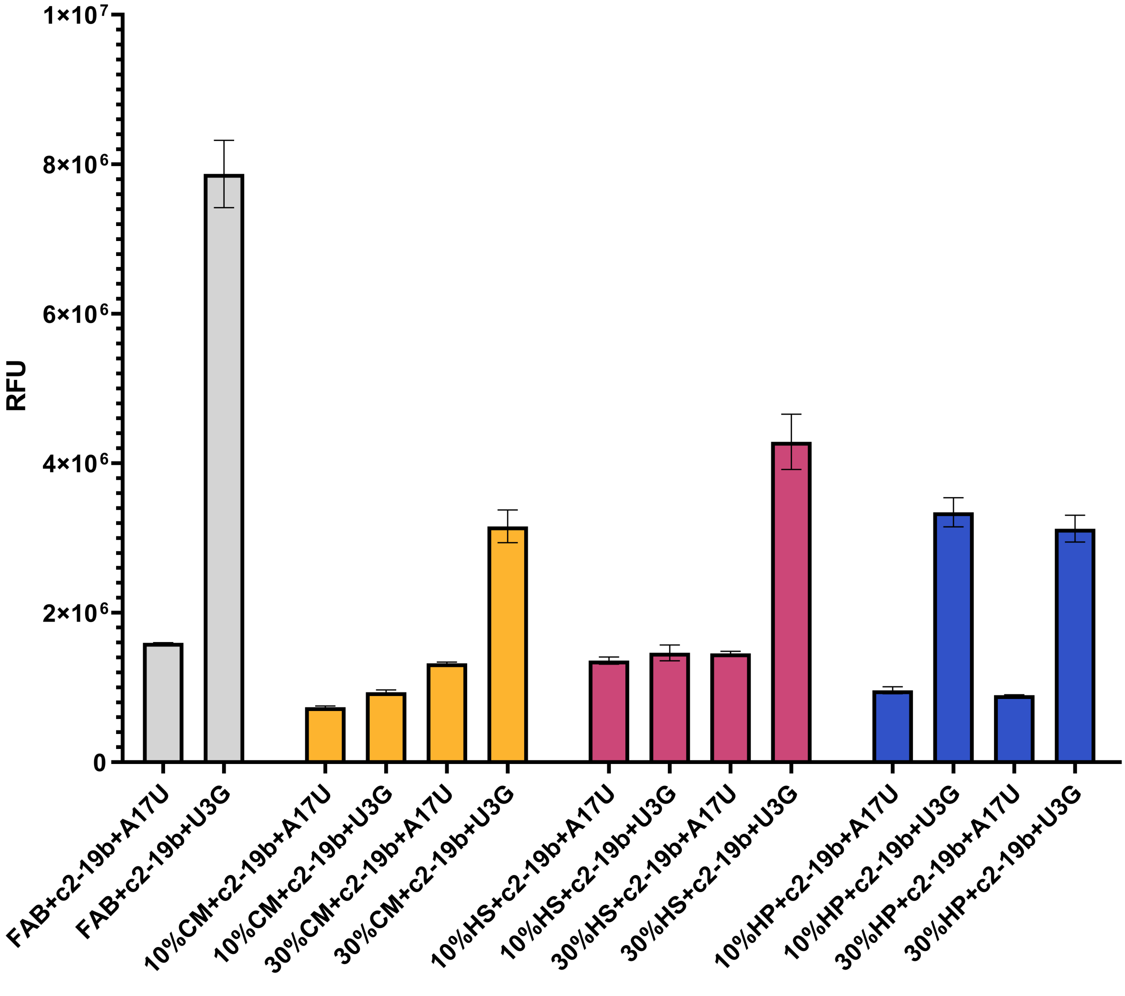

**Supplementary Fig. 5a.** Selectivity of c2-19b against miR-19b mutants A17U and U3G.

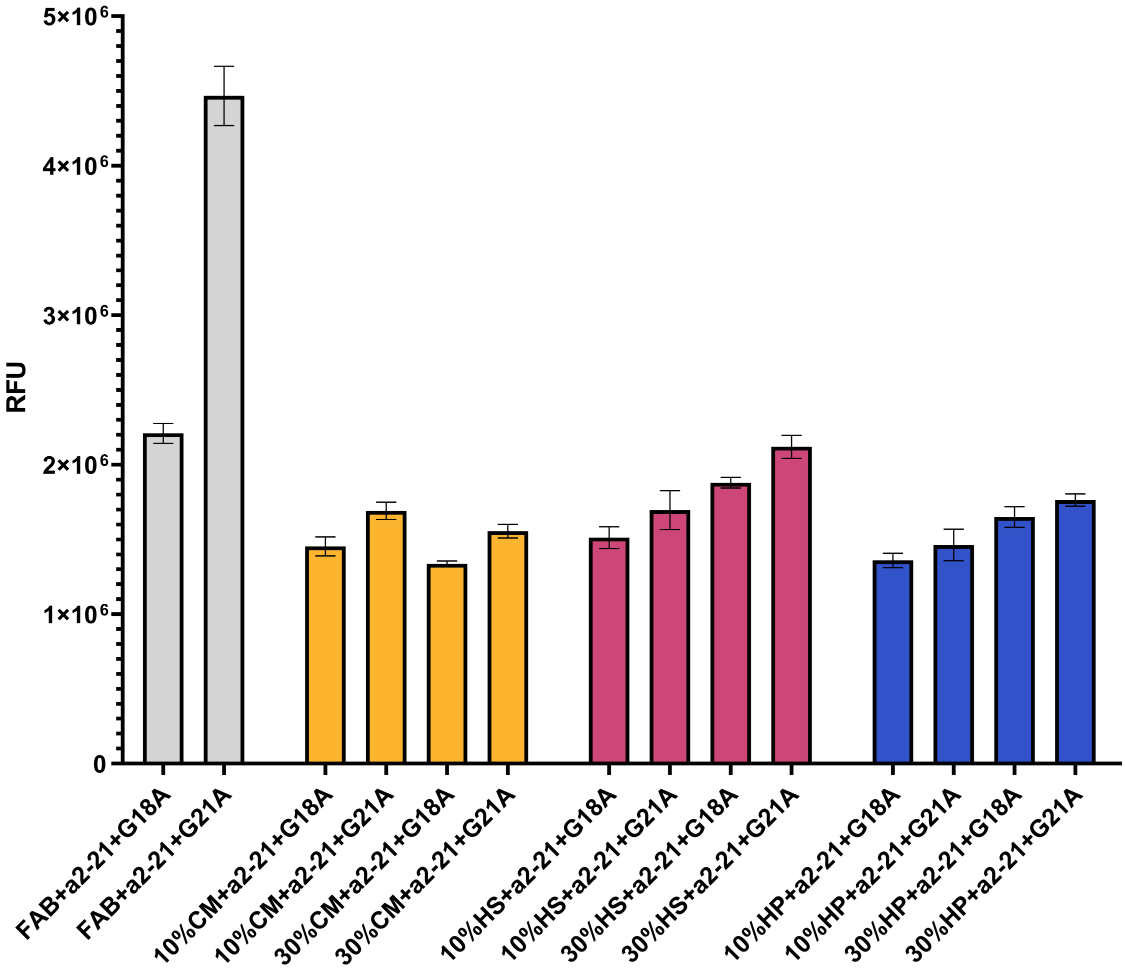

**Supplementary Fig. 5b.** Selectivity of a2-21 against miR-21 mutants G18A and G21A.

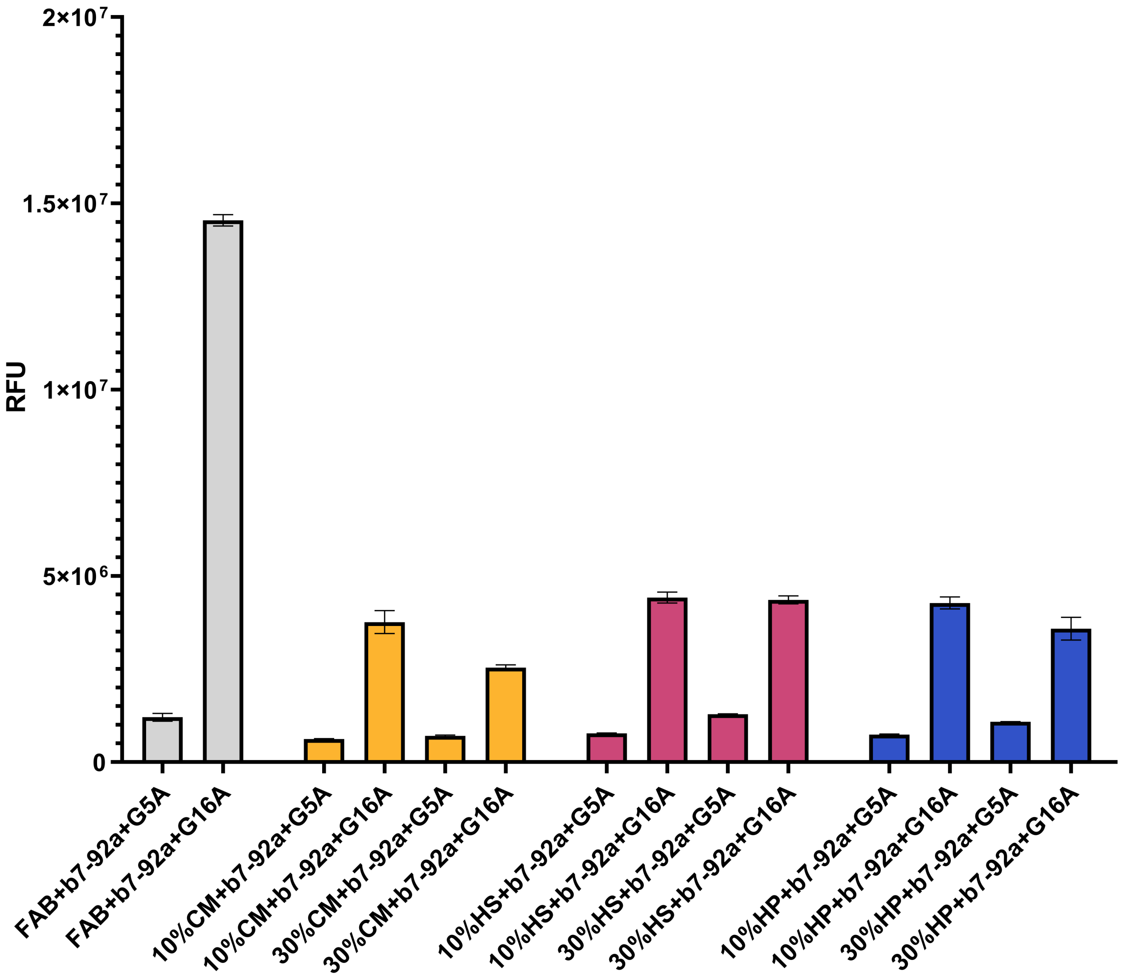

**Supplementary Fig. 5c.** Selectivity of b7-92a against miR-92a mutants G5A and G16A.

**Supplementary Table 5.** Summary of $D_f$ in FAB and biological media.

| miRNA | mutants | FAB | 10% CM | 30% CM | 10% HS | 30% HS | 10% HP | 30% HP |
|---|---|---|---|---|---|---|---|---|
| miR-19b | A17U | 9.8 | 7.1 | 10.3 | 8.2 | 14.6 | 13.7 | 17.9 |
| | U3G | 2.0 | 2.9 | 9.5 | 2.8 | 4.2 | 3.9 | 14.1 |
| miR-21 | G18A | 10.2 | 7.3 | 12.4 | 7.4 | 12.0 | 6.6 | 12.3 |
| | G21A | 5.1 | 6.3 | 11.1 | 6.6 | 11.1 | 6.2 | 10.6 |
| miR-92a | G5A | 23.8 | 18.3 | 18.4 | 13.0 | 19.0 | 11.2 | 24.8 |
| | G16A | 2.0 | 14.3 | 13.3 | 10.7 | 14.9 | 9.7 | 15.0 |

### Limit of Detection (LoD)

LoDs were determined by measuring the fluorescence intensities of samples of AO with St1:St2:miRNA prepared in pure buffer or 10% or 30% (v/v) biological media. Samples contained different concentrations of the target miRNA (0–100 nM for assays in FAB or 0–20 nM for assays in biological media), 100 nM St1, 100 nM St2, and 2 μM AO. Samples were prepared and the fluorescence intensities were measured after 1 hour of incubation at RT. Three replicates of experiments were performed and one-tailed Student's *t*-test was used to assess the statistical difference between the intensity at each concentration of miRNA and that at 0 nM miRNA.
